# Annotation Vocabulary (Might Be) All You Need

**DOI:** 10.1101/2024.07.30.605924

**Authors:** Logan Hallee, Niko Rafailidis, Colin Horger, David Hong, Jason P. Gleghorn

## Abstract

Protein Language Models (pLMs) have revolutionized the computational modeling of protein systems, building numerical embeddings that are centered around structural features. To enhance the breadth of biochemically relevant properties available in protein embeddings, we engineered the *Annotation Vocabulary*, a transformer readable language of protein properties defined by structured ontologies. We trained *Annotation Transformers* (AT) from the ground up to recover masked protein property inputs without reference to amino acid sequences, building a new numerical feature space on protein descriptions alone. We leverage AT representations in various model architectures, for both protein representation and generation. To showcase the merit of Annotation Vocabulary integration, we performed 515 diverse downstream experiments. Using a novel loss function and only $3 in commercial compute, our premier representation model CAMP produces state-of-the-art embeddings for five out of 15 common datasets with competitive performance on the rest; highlighting the computational efficiency of latent space curation with Annotation Vocabulary. To standardize the comparison of *de novo* generated protein sequences, we suggest a new sequence alignment-based score that is more flexible and biologically relevant than traditional language modeling metrics. Our generative model, GSM, produces high alignment scores from annotation-only prompts with a BERT-like generation scheme. Of particular note, many GSM hallucinations return statistically significant BLAST hits, where enrichment analysis shows properties matching the annotation prompt even when the ground truth has *low* sequence identity to the *entire* training set. Overall, the Annotation Vocabulary toolbox presents a promising pathway to replace traditional tokens with members of ontologies and knowledge graphs, enhancing transformer models in specific domains. The concise, accurate, and efficient descriptions of proteins by the Annotation Vocabulary offers a novel way to build numerical representations of proteins for protein annotation and design.

## Introduction

The evolutionary optimization of proteins is achieved through incremental, seemingly random changes to a genetic code that are mostly detrimental implying that the natural protein landscape is hardly exhaustive (1, 2). Even within the space of natural proteins, high-throughput sequencing technologies have far outpaced our ability to characterize genetic constructs (3). Far less than 1% of documented protein sequences have ever been synthesized, let alone annotated (4–6). The immense challenge of exploring biological sequences with extremely sparse data places protein design and annotation as ubiquitous problems in the life sciences. Understanding this vast landscape of proteins is important for studying and treating diseases, as well as elucidating fundamental biology (7). Beyond biological systems, protein design harbors potential in generating sequences capable of valuable tasks, including plastics degradation and recycling, carbon capture and storage, and the generation of novel materials (8–10). While potential applications are numerous and significant, experimental characterization is time-intensive and expensive, heavily limiting the rate of progress as well as training data availability for computational methods. Therefore, there is a vital need for reliable computational methodologies that can translate between sequence and function based on sparse labeled data.

Both protein annotation and design have been a primary focus of the Protein Language Model (pLM) community, where protein sequences are modeled as a semantic language by amino acids, codons, nucleotides, or atoms (11–16). By leveraging large-scale semi-supervised denoising and transfer learning, transformer neural networks have showcased adept numerical representations that correlate to downstream tasks without any labels at all (11, 17). Of interest in biomedical communities, tasks such as Protein-Protein Interaction (PPI) and function prediction were improved with this approach (12, 17, 18). Generating natural seeming sequences from noise has also been possible with pLMs (19–21). However, a more recent study of pLM pretraining strategies suggests that Masked Language Modeling (MLM) is particularly effective for structure-based modeling, injecting many structurally correlated patterns into the pLM latent space (22). Whereas this gives insight into the success of protein folding models (21, 23–29)), we assume that the optimal latent space for annotation should more closely correlate with more abstract concepts like “protein function” and “biological process.” We also surmise that generating proteins for specific properties can be actualized by a closer relationship between a “property” latent space and sequence latent space.

Others have overcome the pretraining pitfalls of MLM over amino acids by applying additional *labeled* contrastive learning to pretrained models. By identifying similar protein pairs, dissimilar pairs, or building triplet datasets, projects like ProteinVec have greatly increased the functional relevance of downstream pLM fixed-length vector or full-token matrix representations (30–32). This has led to excellent protein representation qualities, which enables protein annotation through supervised learning or vector search (30–32). However, approaches that contrast sequences directly require similarity heuristics which impose human bias, defining what sequences or characteristics are inherently similar. One way around this is to assume sequences and their descriptions should inhabit the same embedding space. We observe this in projects, such as ProteinDT, correlating the pLM latent space directly with researcher-deposited natural language embeddings using contrastive learning (33). While this is a promising avenue for *de novo* protein design from prompts, we suspect that natural language is not an optimal interface to the protein language.

We postulate that much of the challenge involved in enabling effective protein annotation and design lies within the inadequacies in our descriptions of proteins, which are highly complex molecules operating under multiscale constraints. For example, natural proteins are optimized around countless considerations including cellular economics (expression energy and pathway efficiency), regulatory mechanisms (allosteric sites, feedback loops, post-transcriptional/translational modifications), and protein lifecycles (chaperone folding, complex formation, proteolysis) (34–40). Most of these qualities are rarely or never mentioned in deposited natural language descriptions. While it is possible to use Large Language Models (LLMs) to format ontology-based annotations to natural language, it requires nontrivial compute and runs the risk of hallucination (41). Despite these challenges, approaches like Mol-instructions have made great strides toward descriptive, machine-readable prompts for molecular design and annotation (42). Here, we ask: Why not just use the annotations as a direct input? A separate vocabulary of annotations.

To work toward descriptive protein property representations that enable the bidirectional translation of sequence and function, we engineered a new tool called the **Annotation Vocabulary**. The Annotation Vocabulary is a collection of human-labeled protein-related ontologies that concisely and accurately describe protein properties. By mapping Enzyme Commission numbers (EC), Gene Ontologies (GO), Interpro domains, and Gene3D domains to a set of unique integers, the Annotation Vocabulary was able to be modeled with transformer neural networks through token embedding. This eliminated the need for similarity heuristics for comparison between sequences by assuming a fundamental relationship between a sequence and its own annotations. Additionally, unlike natural language descriptions posited by researchers, specific properties were described in a consistent way. A transition away from natural language also removed artifacts like filler words, which saved on computation and increased interpretability. Using this vocabulary, we trained various model architectures to leverage protein annotation representations, including:

- **Annotation Transformer** (AT): A transformer network that uses the Annotation Vocabulary to build functionally relevant representations of annotations,
- **Contrastive Annotation Model for Proteins** (CAMP): Leverages AT to curate sequence representations with contrastive learning using a novel loss,
- **Annotation Sequence Model** (ASM): Utilizes a dual vocabulary of sequences and annotations to curate sequence representations with self-attention,
- **Generation Sequence Model** (GSM): Leverages AT to generate sequences from annotation prompts with cross-attention.

CAMP and ASM were evaluated on protein annotation tasks with downstream supervised learning and vector search, demonstrating a high correlation with valuable tasks. CAMP produced SOTA embeddings for five out of 15 standardized datasets with competitive performance on the rest, significantly outperforming the newest foundation model ESM3. To compare the Annotation Vocabulary to other strategies, we compiled a dataset of natural language descriptions and proteins that were applied to CAMP, replacing the AT with SciBERT, which also outperformed pretrained pLMs. Notably, training our premier representation model CAMP_*EXP*_ cost a **total of $3 in commercial compute** (3 hrs on an A6000), highlighting the computational efficiency of latent space curation with Annotation Vocabulary. We conducted protein annotation and sequence reconstruction tasks on AT and ASM using mask filling, both with and without reference amino acid sequences. F1 scores and loss values for sequence reconstruction show that ASM35 can outperform ESM2-150, underscoring added value in incorporating the Annotation Vocabulary into standard pLM pretraining practices.

However, standard metrics like accuracy or F1 scores between reconstructions and labels, as well as loss or perplexity, require the indices of correct tokens to exactly match, which is less meaningful for *de novo* protein generation. Within the context of biological sequences, many conserved domains may function correctly if slightly out of frame meaning a high-quality generation result similar to the ground truth sequence may present poor metrics. For a more standardized comparison of generated biological sequences, we propose a novel normalized sequence alignment score based on the Needleman-Wunsch algorithm (43). Using this metric, we explored how well GSM can generate sequences at various mask percentages with annotation prompts, including from pure noise. Importantly, GSM generated realistic protein sequences with high sequence alignment scores to ground truth. Following Basic Local Alignment Search Tool (BLAST) queries, we show statistically significant hits with sequences annotated similar to the prompted annotations <u>even when the ground truth has a low sequence identity to the training set.</u> Overall, our work offers a new way to build numerical descriptions of proteins through the Annotation Vocabulary. When utilizing our strategies, the functional relevance of amino acid embeddings is enhanced, hinting at broader improvements in both protein annotation and design.

## Results

We used the Annotation Vocabulary to curate the latent space of various transformer architectures. To evaluate the effectiveness of Annotation Vocabulary integration, we performed **515** diverse performance evaluations. This included protein annotation using supervised learning with model probes, vector search, mask filling, and protein design using annotation vocabulary prompts. Throughout, we will refer to *sequence reconstruction*, where we are measuring the capabilities of models to exactly replicate ground truth sequences, and *sequence generation*, where we measure performance with less strict, but more biologically relevant, alignment-based metrics to explore the generation of plausible protein domains.

### Annotation Vocabulary enhances the value of protein embeddings

Firstly, we set out to improve representation learning schemes with the Annotation Vocabulary. We compiled the EXP (UniProt sequences and experimentally validated redundant annotations, 70,000 total), RED (UniRef90 sequences and nonredundant annotations, 500,000 total), and NAT (UniRef50 sequences and nonredundant natural language descriptions, 1.4 million total) datasets. To conduct representation learning over pure annotations, we trained the Annotation Transformer (AT) (**Figure 1A**), a BERT-like (44) transformer, on the EXP and RED datasets separately, named AT_*EXP*_ and AT_*RED*_ respectively. Then, CAMP models were trained with AT components and ESM2-650 (45–50) to curate the ESM2 latent space with annotations through contrastive learning (**Figure 1B**), named CAMP_*EXP*_ and CAMP_*RED*_ respectively. For comparison against natural language descriptions, AT was replaced with SciBERT (51) on the NAT dataset, producing CAMP_*NAT*_. Lastly, ASM (EXP and RED) (**Figure 1C**) has joint representation and reconstruction capabilities, so ASM was evaluated for sequence-only representation as well.

**Fig. 1.**
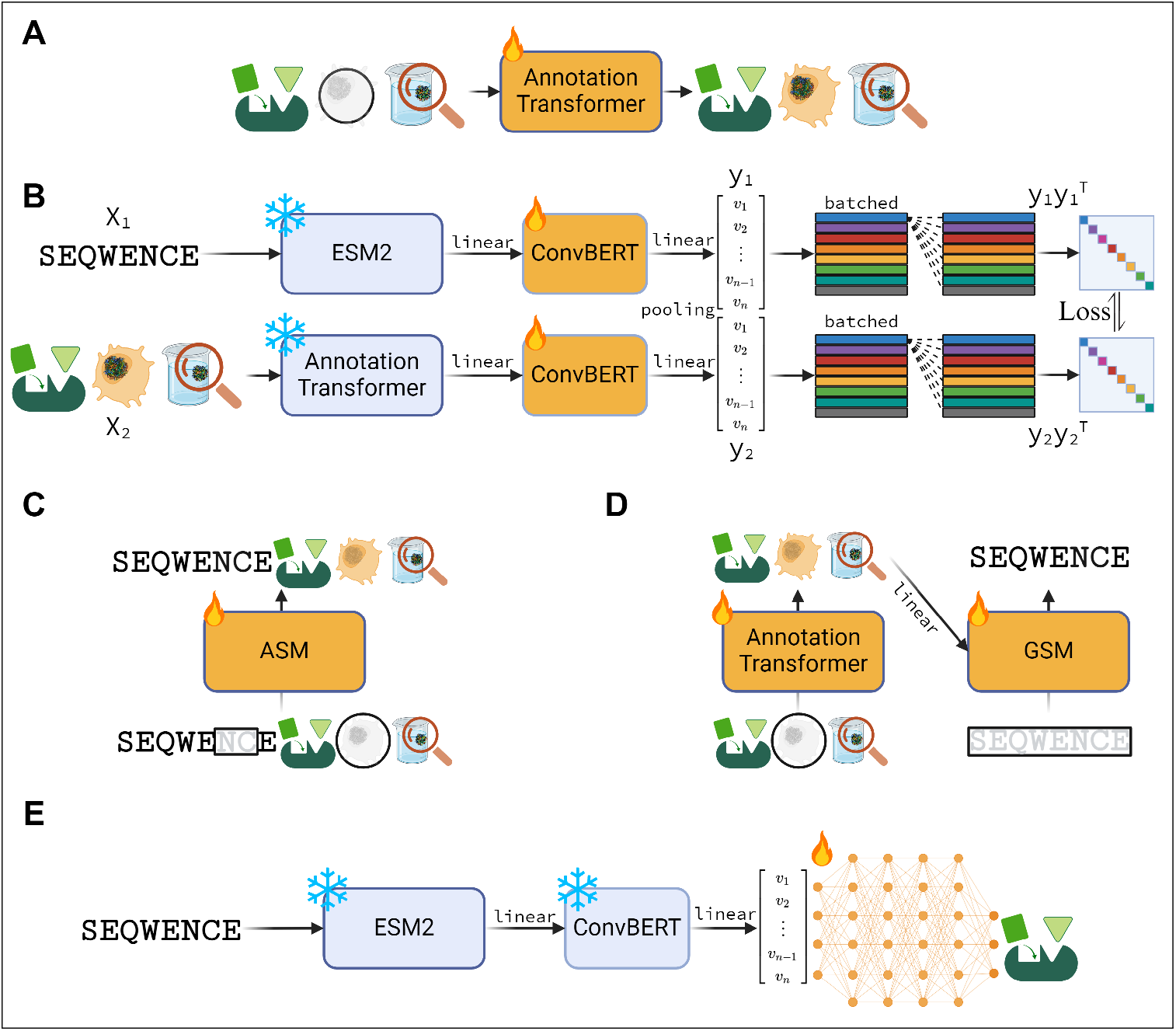
**A:** Annotation Transformer, a BERT-like network trained through MLM on protein annotation tokens. **B:** CAMP model schema, where ESM2-650 and AT are frozen to produce consistent representations after pretraining. Linear layers and ConvBERTs project these representations to a common hidden dimension. Vector outputs are contrasted at the mini-batch level to curate the protein latent space for annotation tasks. **C:** ASM, an ESM model with an extended dual sequence-annotation vocabulary, trained through MLM on both vocabularies. **D:** GSM, where AT hidden states are attended with an ESM2 model through cross-attention to enable protein sequence generation from annotation prompts. **E:** Example of model probe pipeline, where only the sequence track is used and frozen to produce embeddings, then used to train a probe. Created with BioRender.com

We evaluated sequence embeddings after contrastive learning on various tasks, split into in-distribution and out-of-distribution, which were either discretely defined within the Annotation Vocabulary (in) or not (out). Fixed-length vector embeddings from frozen models were fed to a linear probe (**Figure 1E**). As expected, consistent performance increases versus CAMP’s base model ESM2-650 were seen on in-distribution tasks (**Table 1**). CAMP variants exhibited the best overall performance of the tested models, with CAMP_*EXP*_ embeddings resulting in the only average F1 score above 0.6. CAMP_*EXP*_ scores were 2.6% higher than ESM3 (52) and 9.5% higher than ProteinVec, two large models that were trained with functional information on top of amino-acid based pretraining. Individually, CAMP_*EXP*_ embeddings produced the highest DL10 and second-highest EC and CC F1 scores, with CAMP_*RED*_ generating the best CC and BP F1 scores. ASM embeddings also performed well, with RED and EXP embeddings 2.2% and 3.3% higher than their base model ESM2-35 F1 score on average. Interestingly, many smaller models trained by semi-supervised denoising had embeddings that correlated better with downstream tasks compared to larger counterparts. ESM2-150 is particularly good at DL2 prediction, and the best overall non-CAMP model was ANKH_*base*_ (17); matching CAMP_*NAT*_ in average performance with an F1 average of 0.589.

**Table 1.**
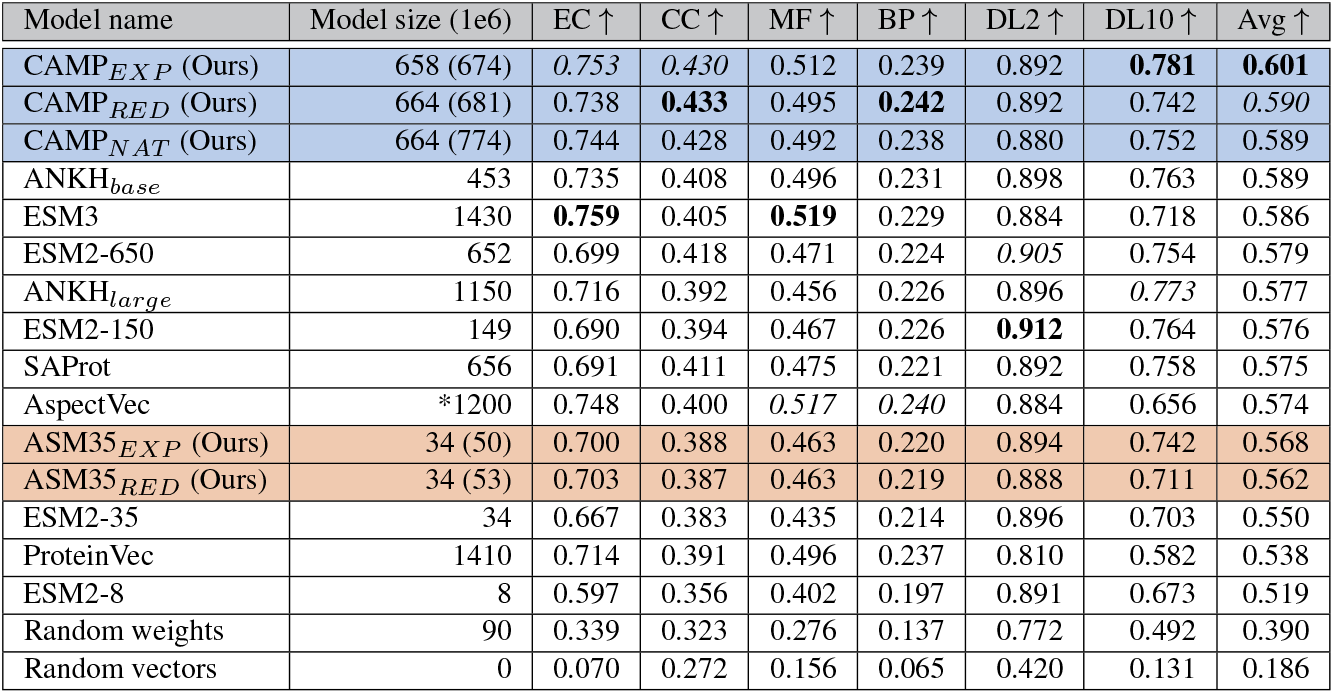
F1_*max*_ (multi-label) and F1 scores shown for *in-distribution* downstream tasks, which we classify as well aligned with the properties represented in the Annotation Vocabulary. Our models have the total model size in parenthesis referencing the total training schema including Annotation Vocabulary components which are not referenced during this embedding process. CAMP model embeddings outperform their closest equivalent counterpart in terms of methodology: ProteinVec, and also the newest frontier pLM ESM3. ASM35 outperforms its base model ESM2-35 in sequence only inference. * approximation of 1 AspectVec on top of ProtT5 encoder (11). EC, CC, MF, BP, CC, and CC AspectVecs refers to the order used in the table (30).

Our out-of-distribution evaluation using vector embeddings and a linear probe portrayed a similar story (**Table 2**). Here, we did not necessarily expect increased performance compared to the base model. CAMP models individually excelled at PPI tasks, scoring the first and second highest F1 for each. However, it is clear that ANKH and ESM variants outperformed CAMP and ASM on MB. The lack of cofactor annotations, including metal cofactors, for RED and EXP is made clear: CAMP_*EXP*_ MB performance is lower than its base ESM2-650, and ASM35_*EXP*_ performed worse than ASM35_*RED*_ even though the EXP variant has sparse cofactor information and RED does not.

**Table 2.** F1_*max*_ (multi-label) and F1 scores shown for *out-of-distribution* downstream tasks, which we classify as outside the scope of the properties represented in the Annotation Vocabulary. While EXP and NAT have sparse CO annotations, RED has none, which is why we classify MB as out-of-distribution. CAMP embeddings showcase adept performance in PPI despite not being trained for it, vastly outperforming other SOTA pLMs with competitive performance in the MB task.

| Model | MB↑ | YPPI↑ | HPPI↑ | Avg↑ |
| --- | --- | --- | --- | --- |
| ESM2-650 | 0.705 | 0.773 | 0.762 | **0.747** |
| CAMP<sub>NAT</sub> | 0.669 | **0.794** | 0.763 | 0.742 |
| CAMP<sub>RED</sub> | 0.646 | 0.782 | **0.767** | 0.732 |
| ANKH<sub>large</sub> | 0.723 | 0.757 | 0.715 | 0.732 |
| ANKH<sub>base</sub> | **0.748** | 0.755 | 0.689 | 0.731 |
| ESM3 | 0.715 | 0.756 | 0.722 | 0.731 |
| ESM2-150 | 0.690 | 0.754 | 0.750 | 0.731 |
| CAMP<sub>EXP</sub> | 0.697 | 0.787 | 0.703 | 0.729 |
| ESM2-35 | 0.691 | 0.745 | 0.732 | 0.723 |
| ESM2-8 | 0.670 | 0.745 | 0.677 | 0.697 |
| ASM35<sub>EXP</sub> | 0.642 | 0.737 | 0.671 | 0.683 |
| ASM35<sub>RED</sub> | 0.661 | 0.702 | 0.661 | 0.675 |
| ProteinVec | 0.635 | 0.550 | 0.539 | 0.575 |

Supervised learning is not the only avenue for protein annotation; embedding labeled datasets and conducting vector search via vector similarity has also shown promise (30, 32). As such, we evaluated EC annotation using vector search and a SwissProt reference database with maximum separation techniques (32) (**Table 3**, full metrics in **Supplemental Table 1**). For the three benchmark datasets introduced by CLEAN (New, Price, Halogenase) (32), CAMP embeddings performed competitively, with the CAMP_*EXP*_ achieving an average AUC of 0.785. Notably, the CLEAN_*EXP*_ scores for New and Price were within 0.001 AUC of the highest performers, ProteinVec and CLEAN, respectively. ASM underperformed compared to ESM2-35 but still outperformed ESM2-650 on average; there was no consistent size-to-performance trend with the CLEAN benchmark.

**Table 3.** AUC for the CLEAN datasets using the maximum separation method with reference to a Split100 (SwissProt) vector database (32). CAMP and ASM models do not outperform their counterparts, but CAMP_*EXP*_ is within 0.001 AUC of SOTA on the New and Price dataset. Unsurprisingly, ProteinVec and CLEAN still perform excellently around their designed purpose of annotation by vector search (30, 32). * Reported (32)

| Model | New↑ | Price↑ | Halogenase↑ | Avg↑ |
| --- | --- | --- | --- | --- |
| ProteinVec | **0.761** | 0.709 | **0.926** | **0.799** |
| *CLEAN | 0.740 | **0.733** | 0.907 | 0.793 |
| ANKH<sub>base</sub> | 0.744 | 0.699 | 0.921 | 0.788 |
| CAMP<sub>EXP</sub> | 0.760 | 0.732 | 0.863 | 0.785 |
| ANKH<sub>large</sub> | 0.747 | 0.728 | 0.862 | 0.779 |
| ESM2-35 | 0.705 | 0.675 | 0.851 | 0.744 |
| ESM-3 | 0.708 | 0.697 | 0.807 | 0.737 |
| CAMP<sub>RED</sub> | 0.706 | 0.703 | 0.783 | 0.731 |
| CAMP<sub>NAT</sub> | 0.698 | 0.680 | 0.797 | 0.725 |
| ASM35<sub>NAT</sub> | 0.708 | 0.677 | 0.777 | 0.721 |
| ESM2-150 | 0.672 | 0.683 | 0.782 | 0.712 |
| ASM35<sub>EXP</sub> | 0.689 | 0.631 | 0.781 | 0.700 |
| ESM2-650 | 0.686 | 0.589 | 0.810 | 0.695 |
| ESM2-8 | 0.688 | 0.637 | 0.713 | 0.679 |

We also evaluated the residue-wise matrix embeddings for CAMP (**Table 4**), even though we used a loss that was based on vector representations. The goal of this experiment was to determine if a pooled vector-based loss inhibits residue-wise tasks. By far, ANKH_*large*_ and ESM3 embeddings exhibited the best correlation with the SS tasks; however, they struggled with TS comparatively. Despite CAMP_*EXP*_ not achieving the top or second best performance for these residue-wise tasks, it still attained the best overall average at 0.652 F1 with ESM2-150 and ESM2-650 slightly lower. ASM models were close to their base ESM2-35 model on SS but significantly underperformed on TS. Similarly to performance with the CLEAN benchmark, TS results were not correlated with model size, as smaller ESM2 models performed the best. Notably, ESM2-8 outperformed comparatively massive models ESM3 and ANKH_*large*_ on average. Additional recorded metrics for protein annotation via model probes can be seen in **Supplemental Table 2**.

**Table 4.** Spearman *ρ* (TS) and F1 (SS3, SS8) shown for annotation tasks using frozen residue-wise embeddings. All Spearman *ρ* values are highly statistically significant (*p <* 1*e*^−5^). CAMP was trained with vector embeddings in mind but is still the best on average. ASM does not outperform ESM here but has competitive metrics.

| Model | TS↑ | SS3↑ | SS8↑ | Avg↑ |
| --- | --- | --- | --- | --- |
| CAMP<sub>EXP</sub> | 0.632 | 0.724 | 0.601 | **0.652** |
| ESM2-150 | 0.656 | 0.710 | 0.586 | 0.651 |
| ESM2-650 | 0.603 | 0.731 | 0.608 | 0.647 |
| ANKH<sub>base</sub> | 0.597 | 0.730 | 0.604 | 0.644 |
| CAMP<sub>RED</sub> | 0.566 | 0.725 | 0.607 | 0.633 |
| ESM2-35 | **0.663** | 0.678 | 0.543 | 0.628 |
| CAMP<sub>NAT</sub> | 0.531 | 0.723 | 0.592 | 0.615 |
| ESM2-8 | 0.632 | 0.670 | 0.521 | 0.608 |
| ASM35<sub>RED</sub> | 0.620 | 0.669 | 0.529 | 0.606 |
| ASM35<sub>EXP</sub> | 0.581 | 0.668 | 0.527 | 0.592 |
| ANKH<sub>large</sub> | 0.337 | **0.749** | **0.649** | 0.578 |
| ESM3 | 0.217 | 0.733 | 0.628 | 0.526 |

### Protein classification is tractable through bidirectional mask filling

We evaluated the performance of AT and ASM35 models in predicting masked annotations by modeling protein annotation as mask filling. We masked one annotation category completely (e.g. EC, CC, MF, etc.), and the models used the remaining annotations to predict the missing ones. For the ASM models, experiments were performed both with annotation information as an input and with annotation and full sequence inputs (denoted ASM35_*X*_ -Seqs) for additional context. The AT models demonstrated superior performance compared to the ASM35 models across five of six downstream tasks (**Table 5**). AT_*EXP*_ had stronger performance in EC, Interpro, and Gene3D predictions, while AT_*RED*_ excelled in MF and BP predictions. ASM35 models underperformed versus AT models across most tasks, with ASM35_*RED*_ only marginally outperforming AT_*RED*_ on the BP task by 0.002 F1 score. However, the addition of sequence information to ASM did improve its F1 scores on average.

**Table 5.** Protein annotation as mask-filling with Annotation Vocabulary. F1 scores are shown for each downstream task. Models were evaluated on their respective validation datasets to fill in a missing aspect given the other. ASM models with “-Seqs” also had the full amino acid sequence as context.

|  | EC↑ | MF↑ | BP↑ | CC↑ | Pfam↑ | Gene3D↑ | Avg↑ |
| --- | --- | --- | --- | --- | --- | --- | --- |
| AT<sub>RED</sub> | 0.633 | **0.498** | 0.543 | **0.392** | 0.352 | 0.527 | **0.491** |
| AT<sub>EXP</sub> | **0.716** | 0.391 | 0.179 | 0.319 | **0.552** | **0.672** | 0.472 |
| ASM35<sub>RED</sub>-Seqs | 0.532 | 0.452 | 0.540 | 0.367 | 0.326 | 0.482 | 0.450 |
| ASM35<sub>RED</sub> | 0.515 | 0.457 | **0.545** | 0.366 | 0.319 | 0.468 | 0.445 |
| ASM35<sub>EXP</sub>-Seqs | 0.153 | 0.143 | 0.032 | 0.165 | 0.140 | 0.327 | 0.160 |
| ASM35<sub>EXP</sub> | 0.135 | 0.145 | 0.031 | 0.151 | 0.132 | 0.293 | 0.148 |

### Sequence reconstruction is improved with annotation context

Next, we set out to establish protein reconstruction schemes leveraging the Annotation Vocabulary. After training, ASM exhibited higher sequence recovery rates by leveraging annotations, highlighting its ability to fill masked regions given annotation context. Examining

ASM35_*RED*_ throughout training (early, mid, late, RED), we observed a gradual increase in reconstruction proficiency (**Figure 2A**) as compared to its base model ESM2-35. While this is true for all mask percentages compared to ESM2-35, ASM35_*RED*_ was much better at high percentage mask recovery even relative to ESM2-150 with 0.23 vs. 0.21 F1 for 50% masking and 0.08 vs. 0.03 F1 for 70% masking. For ASM35_*EXP*_, this difference is even more pronounced, with ASM35 overtaking ESM2-150 on all percentages (**Figure 2C**), including 0.24 compared to 0.23 for 50% mask and 0.08 compared to 0.03 for 70% masking. Even at low corruption rates, ASM35_*EXP*_ outperformed ESM2-35 for sequence recovery (0.39 versus 0.30 F1 at 5%). The loss exhibited similar trends for both ASM35 models (**Figure 2B,D**), revealing that the improvement exists at the logit level and is not only performance improved by exact match recovery. With ASM35 exhibiting significant improvement in reconstruction by leveraging annotation context (even surpassing the much larger ESM2-150), the possibility may exist to extend the vocabulary of existing SOTA pLMs with this Annotation Vocabulary to train further for sequence reconstruction. This may be particularly valuable for tasks such as active site optimization and mutagenesis study (48, 53).

**Fig. 2.**
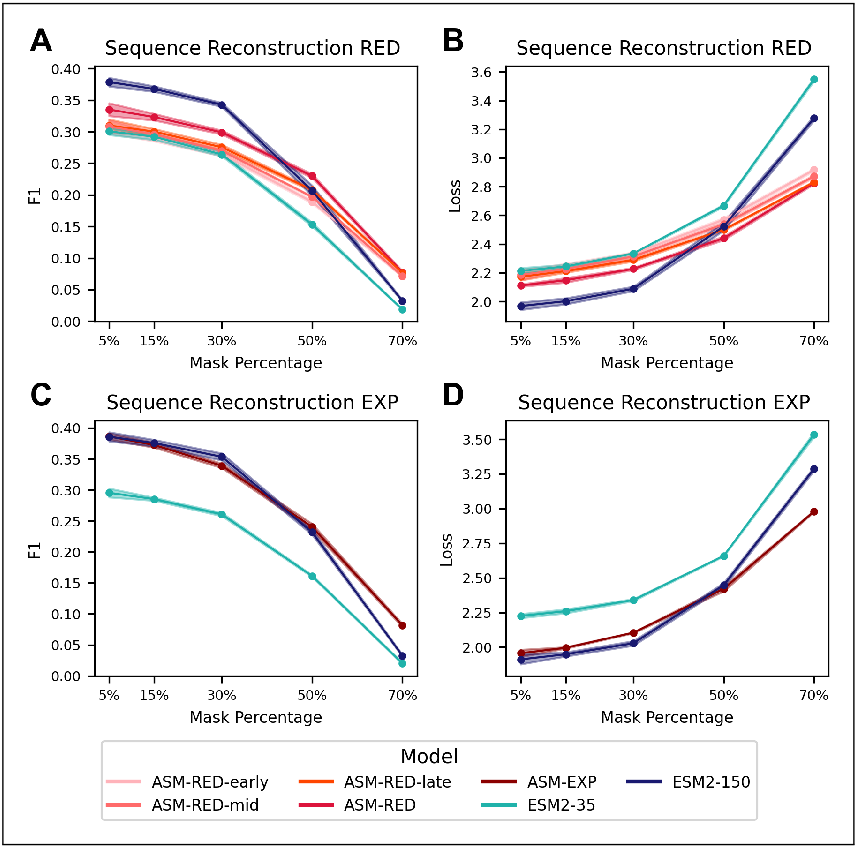
Average performance of sequence reconstruction with three standard deviation error bars. ASM35_*RED*_ models are colored light to dark based on training progress - highlighting improvements throughout training. **A:** Sequence reconstruction F1(↑) of ASM35_*RED*_ sampled throughout training vs. ESM2-35 and ESM2-150. **B:** Sequence reconstruction loss(↓) of ASM35_*RED*_ sampled throughout training vs. ESM2-35 and ESM2-150. **C:** Sequence reconstruction F1 of ASM35_*EXP*_ vs. ESM2-35 and ESM2-150. **D:** Sequence reconstruction loss of ASM35_*EXP*_ VS.ESM2-35 and ESM2-150.

### Annotation Vocabulary prompts generate sequences *de novo* that align well with ground truth

We engineered transformer components to create GSM, which leverages AT to generate sequences from annotation prompts. To evaluate the BERT-like generation performance of GSM against ESM2, we used a novel sequence alignment metric based on the Needleman-Wunsch algorithm and BLOSUM62 to compare how well a generated output matches the ground truth based on evolutionary log-odds. Our metric has advantages in the context of ***de novo*** design compared to traditional metrics like perplexity. Firstly, the metric is scaled from zero (extremely poor alignment) to one (perfect one-to-one amino acid match), which is convenient for interpretation, although it requires more and more similarity to get closer and closer to one. Secondly, it does not penalize models when they move conserved domains out of frame, even though they resemble ground truth almost perfectly. Traditional language modeling metrics look for the exact match in indices, whereas alignment-based methods can introduce gaps to align sequences optimally. Using this metric, randomly paired or generated sequences have a mean score close to 0.15, whereas above 0.5 implies an extremely high degree of similarity. Additional plots to understand possible alignment score distributions are shown in **Supplemental Figure 1**.

In addition to our novel alignment score, we used BLAST to query generated sequences against a nonredundant version of SwissProt (54). When sequences return statistically significant results it indicates that a sequence has conserved domains that, at least partially, resemble natural sequences. By running Blast2GO on BLAST hits, we were able to get high-quality GO annotations through enrichment analysis backed by the manual annotation of SwissProt. This approach allowed us to look for matching properties between the annotation prompt and the Blast2GO consensus.

Before evaluation, we tuned generation hyperparameters for nucleus (*p*) or top-*k* sampling (*k*), as well as the number of tokens to generate each forward pass (*u*) and the temperature (*t*). We found that greedy denoising with *u* = 1 leads to the best score on average; however, higher *k* values (3, 5, 10) and lower *p* reduced the tendency of self-reinforcing repetitions. We observed that this repetition prevention improved the qualitative properties of generated sequences without improving quantitative metrics on average. We also found that *t <* 1.0 led to favorable results in top-*k* or nucleus sampling, but below 0.1 was detrimental. In an effort to maximize performance and reduce computational time, we chose *u* = 10 and greedy denoising for reported metrics. Full results our hyperparameter search are shown **Supplemental Tables 3, 4**.

Following hyperparameter tuning, we fed 1000 random sequences from the GSM train and test set with various masking percentages to GSM and ESM2-150. GSM also received a full annotation prompt. Due to the presence of self-reinforcing repetitions common to these models, we “filtered” results by employing a *χ*^2^ (Chi-square) test over amino acid distributions to reject poorly generated sequences with repetitive regions (55). Below 50%, both models perform markedly better than random mask filling (**Figure 3**, train results in **Supplemental Figure 2**). In fact, every pairwise comparison between unfiltered results is highly statistically significant (*p <* 0.001) using a two-tailed *t*test. However, at low percentages, ESM2-150 is noticeably better than GSM at mask filling on the train and test sequences, although we note that the ESM2-150 training set (Uniref50) overlaps considerably. On 70% masking and above, GSM is far superior in generating sequences by leveraging the annotation prompts. By examining the number of filtered sequences, we saw that the prevalence of generating low-quality sequences with highly repeated amino acids is similar at low masking percentages for GSM and ESM2-150, although in general more GSM sequences are filtered out. This has been observed in many transformer generation schemes, but our current generation hyperparameter search has not solved this (56). However, we saw that annotation prompts were better utilized at 70% and higher, which is a statistically significant improvement above ESM2-150 and random filling. From analysis of multiple sequence alignments, we note that GSM results were fairly bimodal, generating sequences that either resembled natural sequences and reconstructed obvious conserved domains or it got stuck in self-reinforcing repetition loops where a few tokens were repeated.

**Fig. 3.**
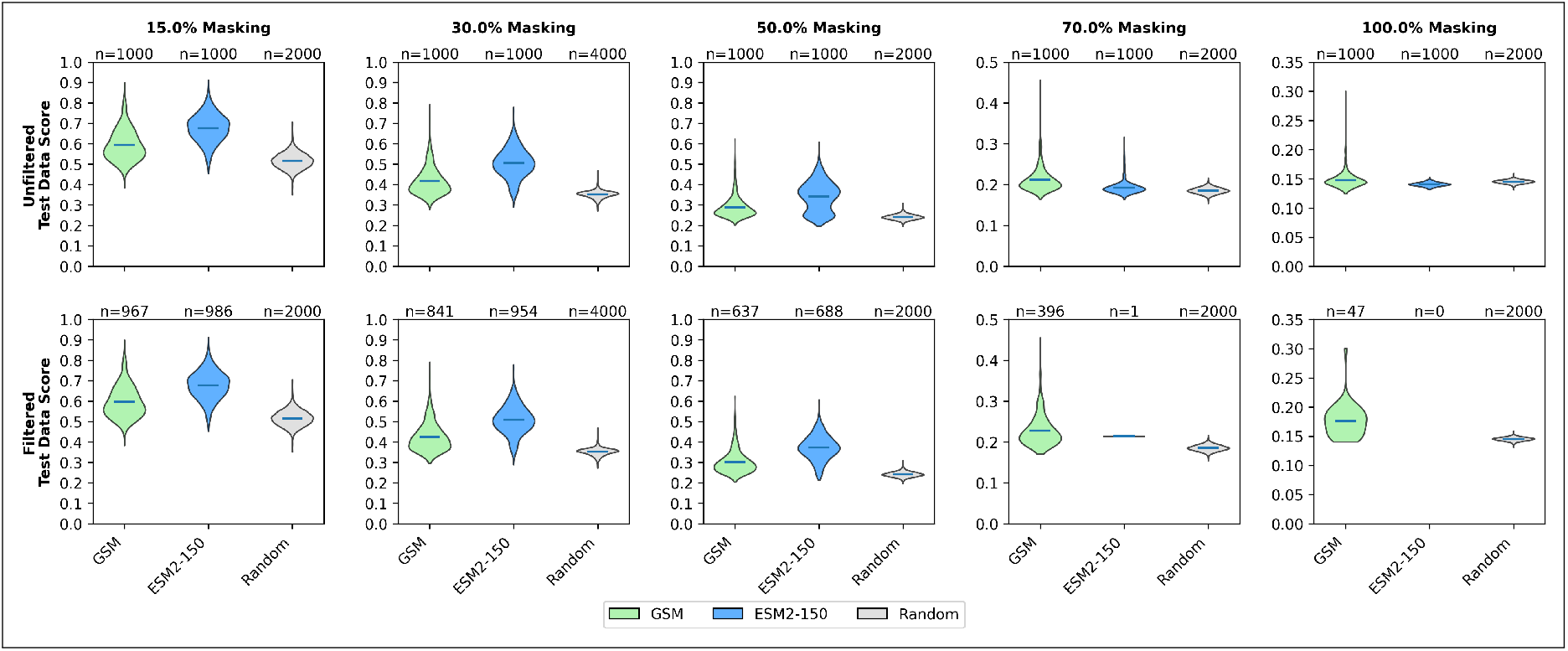
Violin plots of protein sequence generation performance over various mask parameters for 1000 random test sequences. GSM, ESM2-150, and a random mask-filling scheme receive the same masks then fill amino acids with *u* = 10 and *k* = 1 (greedy denoising and 10 tokens at a time). However, GSM also receives an Annotation Vocabulary prompt. Both sets also have low quality results filtered out according to amino acid distribution using *χ*2 test, labeled as “filtered.”

Surprisingly, we do not observe a concrete trend in GSM performance relationship to training set similarity (**Figure 4**). Some test sequences with high training set sequence similarity had near random performance (**Figure 4E**), and some with low sequence identity to the entire training set appeared to be valid proteins exhibiting high alignment scores to their ground truth and returning relevant BLAST hits (**Figure 4B**). GO annotations from all recorded BLAST hits tended to overlap with the input annotation prompt. We suspect that GSM-like models may be trained to condition average results better and that additional schemes such as repetition penalties and MCTS decoding may help (57, 58). Regardless, GSM’s capability of hallucinating natural-like sequences, verified with alignment metrics, BLAST, and enrichment analysis, suggests immense promise in developing generative systems with the Annotation Vocabulary.

**Fig. 4.**
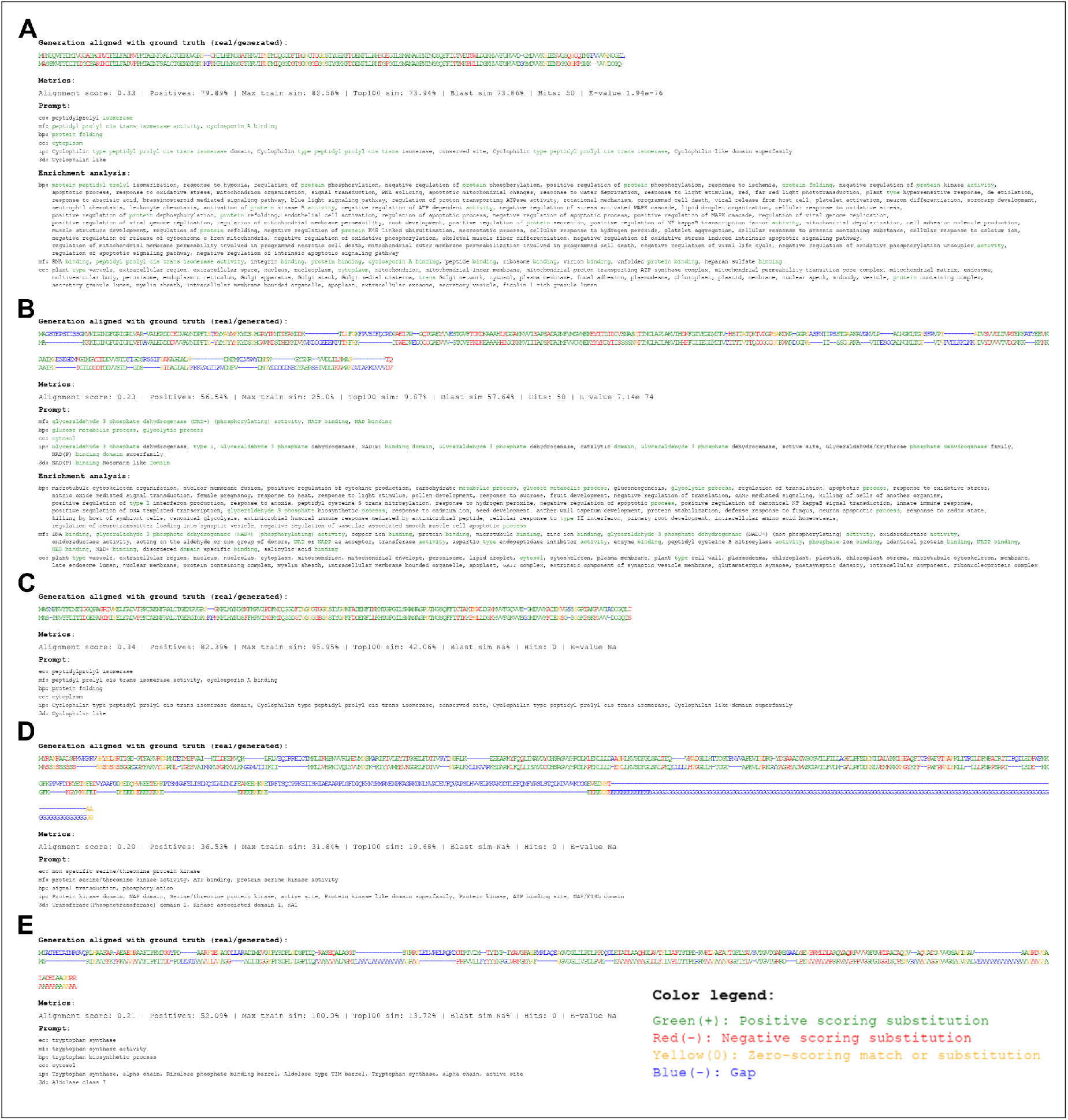
Various GSM sequence generation examples using 100% mask tokens (except a given start methionine) and Annotation Vocabulary prompts (translated to natural language for easier interpretability). Novel alignment score, percent positive alignment indices, max sequence similarity in the training set (Max train sim), the average similarity of the top 100 most similar training sequences (Top100), average sequence similarity of the BLAST hits, the number of BLAST hits, and the BLAST E-value. If a sequence resulted in statistically significant BLAST hits, GO enrichment analysis is shown from Blast2GO (54). Matching words are highlighted in green between the annotation prompt and enrichment terms. **A:** High sequence alignment score (for 100% mask), *high training set similarity*, BLAST hits, matching prompt and GO terms from BLAST hits. **B:** Medium sequence alignment score, *low training set similarity*, BLAST hits, matching prompt and GO terms from BLAST hits. **C:** High sequence alignment score, *high training set similarity*, no BLAST hits. **D:** Medium sequence alignment score, *high training set similarity*, no BLAST hits. This example exhibits highly repetitive regions but also exhibits clearly generated and potentially important domains. **E:** Medium sequence alignment score, *low training set similarity*, no BLAST hits. This example exhibits high repetitive regions but does not exhibit obvious domains.

## Discussion

Recent work suggests that MLM over amino acid sequences instills a representation centered around structural information, where structure-based task performance is disproportionately increased (22). With the goal of annotating sequence repositories on the scale of UniProt, we aimed to curate protein latent spaces that more highly correlate with functional characteristics. To more effectively describe protein annotations at the embedding level, we created the Annotation Vocabulary, a compilation of EC, GO, Interpro, and Gene3D ontologies that map to unique integers. For simplicity, we will refer to unique sets of categorizations (EC, GO, etc.) as aspects, in line with the terminology usage in ProteinVec (30). By assigning token embeddings to each ontology, we hypothesized that we could use annotation tokens to build semantic protein function representations. Our annotation transformer (AT) accomplishes this task, providing a high annotation recovery rate through an MLM objective (**Supplemental Table 5**). When masking out the entirety of specific aspects, AT was able to leverage the other aspects to annotate proteins without reference to sequence information. Interestingly, AT_*EXP*_ had EC prediction F1 score of 0.716 on its validation set without sequence information. This EC annotation task is technically an easier objective than the multi-label task shown via model probes, as the models know how many EC numbers each example should have due to the number of tokens present. Also, some MF tokens or other labels possess a large correlation with EC. However, because this is a normal F1 score and not F1_*max*_, implying this is actually an exceedingly high metric compared to probe-based reported metrics. Sequence annotation as mask filling opens the door to labeling many of the partial annotations in UniProt as our techniques mature, with and without amino acid sequence context.

Once we engineered the Annotation vocabulary toolbox, we identified four main mechanisms in which to add functional information to a standard ℝ^*L×d*^ hidden state from a transformer-like network: 1) by contrasting, constraining, or regularizing a hidden state with functional embeddings, 2) along *L* with new function tokens, 3) along *d* by elongating the hidden dimension with functional embeddings, and 4) within *d* by concatenating a hidden state with functional embeddings.

The first strategy is inclusive of conventional fine-tuning techniques, where contrastive learning is applied with natural language descriptions or between sequences based on similarity heuristics to curate the latent space after pretraining. In initial attempts at this problem, we designed a protein annotation model inspired by ProteinVec and Mixture-of-Experts frameworks, which we call MOESM. While this system had compelling results for EC prediction (second to only ESM3 by 0.002 F1_*max*_) it did not perform well on average (**Supplemental Figure 3 and Table 6**). Importantly, we concluded from this experiment that small models on top of larger pretrained and frozen pLMs could drastically alter the final functional relevance of the output embeddings, implying an effective and computationally efficient shortcut to full model fine-tuning. With this in mind, we used versions of the AT with ESM2-650 to create our CAMP models, which contrast semantic protein and annotation representations. Our novel loss focused on matching the distribution of *sequences compared to other sequences* with the distribution of *annotations compared to other annotations*. A sequence and its corresponding annotation representation were never directly compared, as would be done with a typical cosine similarity or MNR loss (59, 60). Despite this, CAMP model sequence-only inference resulted in fixed-length vector representations that outperform all tested popular pLMs on downstream probes. In particular, on in-distribution annotation tasks, CAMP_*EXP*_ had the only average F1 score above 0.6, higher than premier (and much larger) models ProteinVec and ESM3. CAMP versions performed with much higher metrics on PPI tasks and do not fall short on residue-wise annotation tasks, despite not being trained for either. We hypothesize that CAMP models may be adept at PPI prediction due to their high performance on BP, as many interacting proteins likely fall in the same BP categories. Additionally, CAMP models also produced the best average F1 score on residue-wise downstream tasks, even though they were not trained with a residue-wise objective. This suggests that pooled representations may be sufficient to curate the entire *L × d* hidden state.

Strategy two offers a theoretically sound way to directly move residue embeddings into more functional clusters. The self-attention mechanism is a built-in vector similarity heuristic in the transformer neural network, which models multi-scale relationships between input tokens through projections and dot products. Therefore, if a model can learn to effectively attend discrete sequence and function tokens, their projections must be moved closer within the embedding space. This strategy was prototyped by ASM, where the ESM2-35 vocabulary was extended with the Annotation Vocabulary to model sequences and annotations in a bidirectional manner. After training, the pooled vector embeddings had an increased correlation to in-distribution annotations with 3.3% higher average F1 compared to ESM2-35. Additionally, sequence reconstruction was greatly improved by referencing annotations, outperforming ESM2-35 and ESM2-150 in mask-filling tasks. Of course, one of the main disadvantages of this approach is that the attention mechanism scales *O*(*L*^2^) with combined protein and annotations length *L*, ultimately posing a significant computational expense.

To work against the problematic attention scaling, and to further prototype protein generation using the Annotation Vocabulary, we designed GSM: An Encoder-Decoder schema using AT to produce rich representations of annotation prompts and generate sequences via a cross-attention mechanism. Notably, we used a BERT-like ESM2 model as the decoder, highlighting the newfound potential of using BERT models for sequential generation similar to diffusion models. By removing mask tokens sequentially and strategically, one bridges the gap between representation and generative modeling, spearheaded by ESM3 (52). We evaluated various generation schemes while comparing GSM and ESM2-150 to random mask filling to assess whether Annotation Vocabulary prompts can give an edge over well-trained models such as ESM2. Thus far the most significant approach for improved generation quality is greedy denoising one token at a time. This can be prohibitively expensive for long sequences at *O*(*L*^3^), but we have found that up to 10 tokens at a time reduces this cost without sacrificing much performance. Importantly, the *L* here scales with the amino acid sequence length primarily, as they are much longer as average, due to the use of a cross-attention mechanism instead of self-attention for annotation information mixing.

While GSM underperforms compared to ESM2-150 when generating sequences with mask percentages at or below 50%, we see the significant value of annotation prompts in GSM’s ability to design sequences at 70% masking or from complete noise. Through manual experimentation by prompting from the test set, we observed that GSM generation was fairly “bi-modal,” either designing a sequence that aligns with some domain to the ground truth or getting stuck in self-reinforcing repetition. We removed poor-quality generations using a *χ*^2^ test as a filter. Some GSM-generated outputs returned BLAST hits, and further enrichment analysis found GO annotations matching the prompt. In particular, **Figure 4B** showcases an example where the ground truth sequence has *low* sequence identity with the *entire* training set. This *hallucinated* protein with many BLAST hits and matching enrichment terms implies that the GSM scheme and Annotation Vocabulary are promising avenues for protein design.

Strategy three is compelling if there were adequate residue-wise protein function ontologies. Whereas there are typically Interpro annotations for every sequence in our dataset (90+%), we are reluctant to rely on mappings that correlate sequence motifs directly to protein function, as conceptually, we would rather include Interpro and GO annotations independently to allow the model to learn an (approximately) optimal relationship. That being said, domain-level correlations to function through sequence homology is a remarkably powerful predictor, and clearly Interpro2Go and the excellent tools that have used it to predict GO terms (61, 62) have significant value. Our experiments evaluating ESM2 models with random weights support sequence homology as a significant driver of protein function similarity. While this seems trivial, vector embeddings from randomized ESM2 weights performed much better with probes than random vectors of the same size alone, as shown in our baseline for in-distribution tasks (**Table 1**). We hypothesize this is because similar proteins by homology will be embedded similarly through the token embedding process, even with random weights. Therefore, the downstream probe was still able to recognize functional clusters within sequences clustered by homology. In addition to the conceptual problem with strategy three, there is the less discussed computational scaling of the MLP sections of transformers which scale *O*(*d*^2^) for hidden dimension *d*, implying adding function regions along *d* would add considerable computational cost.

The fourth strategy has recently been tested with ESM3, whereby function embeddings were added directly to sequence (and other modalities) embeddings similar to token type or position embeddings. Computationally, this does not add much cost to the forward pass as the hidden state size is not augmented. Additionally, this has advantages in any-to-any generation due to its ability to represent diverse prompts across modalities as reported in detail in the recent ESM3 paper (52). We hypothesize that ESM3’s functional integration may limit the range of sequence-wise functional ontologies that can be effectively used, as they are still applied at the residue level. However, the strategies we employ may limit the ability of the model to correlate residues with specific functional characteristics because we do not assign them to residues directly. This speculation points to the optimal strategy as potentially being some combination of these approaches. Importantly, there are some limitations to the evaluation approaches used and some surprising results. For example, small linear or BERT-like probes assessed how directly model *embeddings* correlated with a downstream task, but not the propensity for a model to be fine-tuned for a specific task. Because we were analyzing training strategies to incorporate functional information inherently, this was ideal. However, this approach is less ideal for scaling to a production-ready model. This evaluation strategy produced some results that did not follow a conventional size-to-performance ratio. We hypothesize that this is particularly common for models trained through only semi-supervised denoising, where the local minimum the model has found to minimize language modeling cross-entropy just happens to place downstream embeddings in a way that benefits one task over another.

Another surprising result was that ASM performed worse than AT on annotation mask filling, even when ASM had the context of the entire sequence. In the limit, it is clear that sequence information should not hinder a models’ ability to annotate based on other annotations, as it is additional information. In this case, this could be because ASM was under-trained, or perhaps starting from a pretrained ESM2 checkpoint is not ideal for a dual vocabulary scenario. While perhaps less likely, there could also be some percentage of incorrect annotations (5, 6). We see a similar trend, with GSM performing worse than ESM2 with lower mask percentages for design tasks. In theory, more information with annotations should always improve this performance as it provides more information; however, this information may constrain the generation in a harmful way. The high repetition nature of GSM during inference could also be due to under-training for that specific task. Of course, we cannot confirm that any generated result from ASM, ESM, or GSM is “wrong” without experimental validation, but ground truth comparisons seem to be the best computational equivalent. Lastly, it was surprising that the ESM3 embeddings performed worse on average compared to CAMP and ANKH in spite of its impressive training schema. However, it is important to note that ESM3 was not trained solely for representation learning but generation as well, and thus, probing its embeddings is not necessarily indicative of its value as a whole.

Overall, the Annotation Vocabulary toolbox presents a promising pathway to replace traditional tokens with members of ontologies and knowledge graphs, enhancing transformer models in specific domains. We use these strategies to build a language around protein properties, which we feed to various transformer neural network schemes to enhance computational protein design and annotation.

## Methods

### Annotation vocabulary

The Annotation Vocabulary uses EC, GO Cellular Component (CC), GO Molecular Function (MF), GO Biological Process (BP), Interpro, and Gene3D ontologies to describe protein sequences. For each property / aspect / ontology, a minimum and maximum range of integer values was determined based on the number of possible options within the ontology (for that dataset). Each ontology member was assigned a unique integer in ascending value from EC to Gene3D in the order mentioned above. This resulted in a vocabulary of 30,000 - 60,000 unique integers and annotations depending on the base dataset used for this mapping. Once mapped to integers, annotations were fed to transformer neural networks to build numerical representations after token embedding. Importantly, we always fed annotations to transformer models sorted by their tokenized integer value. This introduced annotation “grammar” and enabled training through semi-supervised denoising.

### Data compilation

We compiled three datasets of protein and annotation pairs called EXP, RED, and FINAL for short. Technically, Pfam annotation from UniProt (4) was used instead of Interpro for EXP and RED. The first dataset focused on experimentally validated annotations (EXP) and was gathered from a UniProt query on 5/20/24. We searched for sequences with experimentally validated (manual) GO annotations that also had at least one EC annotation, and Interpro or Gene3D. Whereas cofactor (CO) annotation was sparse, within this query of experimentally validated and high UniProt annotation-score entries, we also recorded CO information for the EXP Annotation Vocabulary. We removed duplicate sequences primarily by prioritizing the most annotations and secondarily by prioritizing the length of the sequence. We removed sequences of less than 50 amino acids or greater than 2048 for computational efficiency. This resulted in a total of 70,395 sequence annotation pairs. 1000 pairs were randomly withheld for validation.

The second dataset was designed to maximize nonredundancy and size (RED), compiled from a UniProt query on 5/29/24 for sequences with any EC annotation, totaling over 41 million. We kept sequences that were representative sequences for a Uniref90 cluster (63). This struck a balance between accurate annotations (representative sequences are chosen based on UniProt annotation-score) and nonredundancy (maximum 90% pairwise sequence identity) and resulted in a total of 17 million sequences. We used 90% clustering instead of 50 or 30 to maximize the size of the final processed dataset. We saved a full set with all 17 million sequences and annotations (RED_*ALL*_), and a set where duplicate annotation entries are removed (RED) consisting of 516,184 pairs. Duplicates were removed by prioritizing sequence length. 1000 pairs were withheld randomly for validation.

The FINAL dataset was simply a combination of the best characteristics we observed from RED and EXP through experimentation. We constructed 700k sequence annotation pairs that were Uniref50 representative sequences (nonredundant) with a maximum length of 512, 157k experimentally validated sequence annotation pairs (redundant) with a maximum length of 512, and 104k experimentally validated sequence annotation pairs (redundant) with length between 512 and 2,048. We also created a set of nonredundant *annotation only* inputs (no matches), which comprised 212k total entries that were used to train AT_*FINAL*_.

To compare our approach versus more common natural language representations, we used a previously compiled dataset from our lab of protein sequences and natural language descriptions using UniProt called NAT for short. It was compiled from Uniref50 representative sequence and property pairs by adding corresponding headers for each unique annotation type, followed by new lines. For example “EC: 1.1.1.1 \n Localization: cytosol, etc.” Representative sequences were recorded when they exhibited at least three of the annotations in **Figure 5**. Because we used Uniref50, sequences have a maximum sequence identity of 50%; this is not necessarily true of the descriptions, which can match exactly. Therefore, we removed duplicates which resulted in 1,435,224 million examples. Sequence overlap between dataset splits can be found in **Supplemental Figure 4**.

**Fig. 5.**
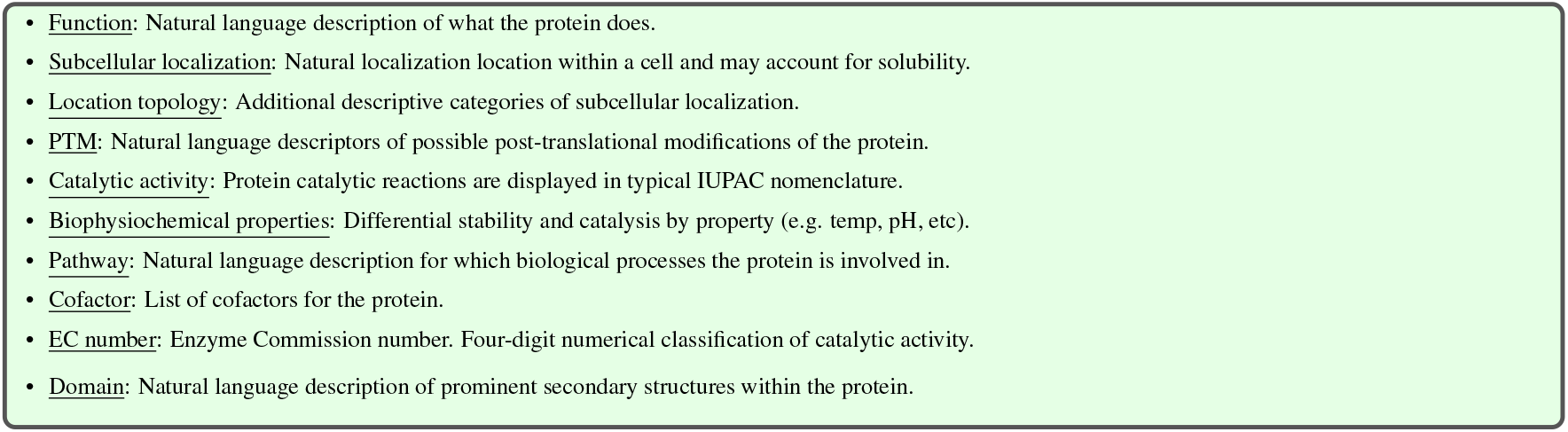
Annotations that defined inclusion criterion for sequence and natural language dataset NAT.

### Annotation Transformer (AT)

We trained two versions of independent AT on EXP and RED, respectively. AT_*EXP*_ is a single BERT-like transformer block with a hidden size 384, intermediate dimension of 2,048, and Annotation Vocabulary of 33,328 (**Figure 1A**). AT_*RED*_ is the same model with an adjusted vocabulary size of 38,953. We also used rotary embeddings instead of absolute position embeddings due to the larger vocabulary size (64). AT_*EXP*_ and AT_*RED*_ were subject to 15% masked language modeling (MLM) objectives for 100 and 10 epochs, respectively, and evaluated periodically based on MLM accuracy on validation sets with early stopping once a patience of three was achieved.

### Annotation Sequence Model (ASM)

The Annotation Sequence Models (ASM) were designed to mix information between sequences and annotations through the self-attention mechanism. These models were actualized by a combination of ESM2 and AT through vocabulary extension - where the token embedding matrix of ESM2 was extended with our Annotation Vocabulary (**Figure 1C**). Like AT, we trained two versions on EXP and RED, respectively. For both experiments, we used ESM2-35 to strike a balance between functional correlation and computational efficiency. Therefore, the two models were called ASM35_*EXP*_ and ASM35_*RED*_ to delineate the dataset they were trained on. Whereas both models were closer to 50 million parameters with the large token embedding matrix and language modeling heads, the weights used during sequence-only inference were exactly equivalent in size to ESM2-35. Both models were subject to 15% MLM objectives with varied training schemes and allowed maximum lengths. When a sequence annotation pair exceeded their combined maximum length, the annotations were shuffled and discarded as the primary truncation strategy to prevent feeding the model protein fragments. However, if this would get rid of over 75% of the annotations, the sequence is truncated as well. ASM35_*RED*_ was first trained on R ED_*ALL*_ for approximately 0.25 epochs (4.25 million sequence annotation pairs) with a maximum length of 768. It was then trained for two epochs on RED with a max length of 1,536. ASM35_*EXP*_ was trained on EXP with a maximum length of 2,048 for 23 epochs total, decided by over-training as observed by decreased MLM recovery accuracy on the validation set.

### Novel contrastive loss

*While directly minimizing the difference between multiple modalities makes the translation between them conceptually convenient, we see no reason to assume that the best latent representation for a protein must be near the representation for its annotation*. Therefore, we designed a novel contrastive loss that instead seeks to match the *intra-latent* relationships among proteins with those of the corresponding annotations. The first term of the loss is the MSE between the pairwise cosine similarities of the protein vector representations and the pairwise cosine similarities of the corresponding annotation representations. This term can be trivially minimized by any solution which projects all outputs to a single point in space, e.g., by zeroing out weights. Hence, we regularize the loss by adding the average intra-latent cosine similarities for each modality to encourage intra-modality embedding diversity. Formally, the loss is defined as follows:

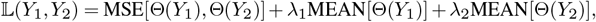

where the rows of *Y*_*i*_ ∈ ℝ ^*n×d*^ are the CAMP outputs for modality *i*, and

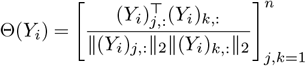

is the corresponding *n* × *n* matrix of pairwise cosine similarities. We chose *λ*_1_ = 1.0 and *λ*_2_ = 0.1 to place emphasis on the diversity of protein representations.

As usual, computing this loss (and its gradients) over the full data is computationally prohibitive, so we instead work in each iteration with stochastic mini-batches 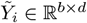 of *b* samples chosen uniformly at random (without replacement) from the full data. Formally, 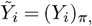 where *π* ∈ {1,…, *n*}^*b*^ denotes the *b* randomly selected sample indices (the same indices are used for all modalities). Notably, the resulting stochastic gradients are technically biased since

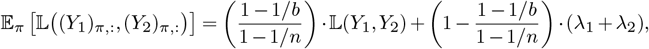

but the resulting scaling can be straightforwardly corrected or even simply ignored. See full details and the derivations on the loss analysis in **Supplemental Loss Analysis** section.

### Contrastive Annotation Modeling of Proteins (CAMP)

The CAMP models are designed to contrast sequence and annotation representations from independent frozen models to further curate the *sequence* latent space (**Figure 1B**). ESM2-650 and AT were frozen, and a small one-block ConvBERT was added to the end of each model alongside many additional linear layers. Sequences were fed to ESM2-650 and corresponding annotations to the AT, which produced full token matrix embeddings. We applied mean pooling to each modality and used our contrastive loss to train the model. CAMP_*EXP*_ and CAMP_*RED*_ delineate which dataset and AT were used. Importantly, the vocabularies are different sizes, so EXP cannot be used to fine-tune CAMP_*RED*_ and vice-versa. For the natural language comparison, we trained CAMP with a frozen SciBERT instead of AT on the NAT dataset. All versions were trained for one epoch over their respective dataset.

### Model benchmarks

To benchmark CAMP and ASM against popular pLMs we designed a rigorous evaluation scheme based on freezing the respective pLM, embedding an entire dataset, and either training a downstream probe or conducting vector search (**Figure 1E**). The datasets used for downstream analysis are shown in **Figure 6**. We used datasets *as is* except for SS and PPI tasks. For SS3 and SS8 we used the Proteinea training set for training (17), CB513 and TS115 for validation (65, 66), leaving CASP12, CASP13, and CASP14 for testing (67). Instead of hiding intrinsically disordered residue labels from the loss function, we created a new label for those residues. Therefore, there were four and nine options per residue for SS3 and SS8, respectively. The PPI sets were the human and yeast splits from PiNUI (68). We generated new validation and test sets using 2500 randomly selected pairs each.

**Fig. 6.**
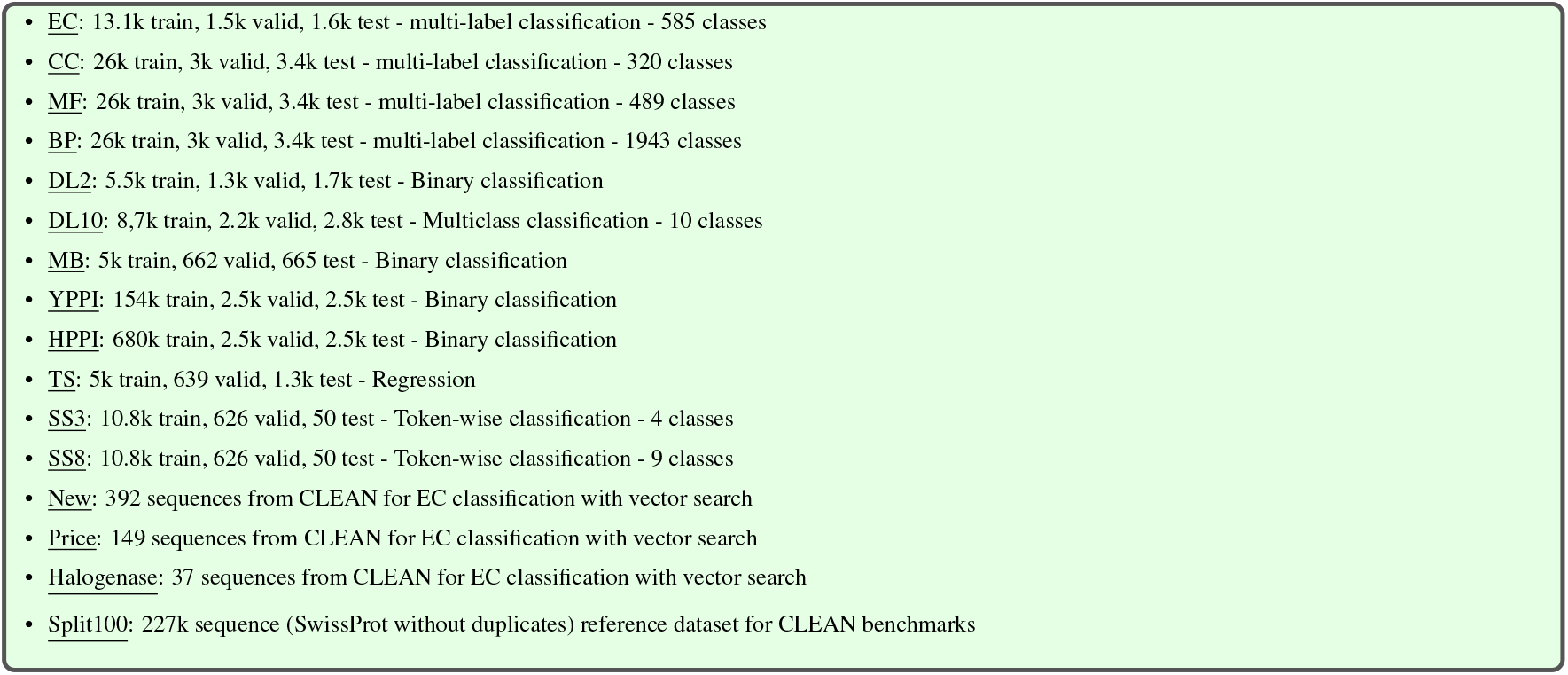
Standard datasets used to probe model performance. EC, CC, MF, BP, DL2, DL10, MB, and TS are from the SaProt project (69). YPPI an HPPI are sampled from the PiNUI project (68). SS3 and SS8 are modified from Protinea (ANKH) (17). New, Price, Halogenase, and Split100 are from the CLEAN project (32).

EC, CC, MF, BP, DL2, DL10, and MB were downloaded from SaProt (69). We note that we fed the amino acid and 3Di sequences to SaProt in the in-distribution benchmark, and just amino acid sequences to the rest (**Table 1**). These datasets, along with YPPI and HPPI, depict sequence-wise categories and were thus evaluated with a linear probe on fixed-length vector embeddings from mean pooling of the last hidden state. For the PPI tasks, we stacked two vector embeddings into a pair representation for each interaction pair. The paired vectors had their order switched with a 50% chance during training. TS is a sequence-wise measurement but is not modelable via a linear probe from a frozen model (all evaluated pLMs perform poorly and inconsistently from pooled states). Therefore, we evaluated TS with a ConvBERT and max pooling, which has been shown to be effective (17, 70). SS3 and SS8 are residue-wise tasks so they were modeled with a BERT probe. The linear probe was a three layer MLP with two hidden layers with a size of 8,192. We tried a large variety of more shallow, more deep, and smaller or larger hidden dimensions and choose this ultimate size based on average performance. For the BERT-like probes, we used an initial linear layer to project the pLM embeddings being evaluated to a standard hidden dimension of 384. We used an intermediate dimension of 1,024 and only used one transformer block for each task. All probes had GELU activation functions.

Probes were trained up to a maximum of 200 epochs to force early stopping, which was triggered by a patience on validation loss improvement, and then evaluated on the test set with the best set of weights. A patience of 10 was used for everything except for SS3 and SS8, which had a more stable performance convergence, and thus we used a patience of five to save time. We validated every epoch except for PPI tasks due to the dataset size, which instead were validated every 1000 batches. A learning rate of 1*e*^−4^ was used for all probes with 100 warm-up steps, a cosine learning rate scheduler, and batch size 64.

CLEAN benchmarks New, Price, and Halogenase were evaluated using the CLEAN maximum separation scheme against an embedding dataset of Split100 from mean pooling (32).

### Protein annotation as mask filling

To evaluate the annotation capabilities of the AT and ASM35 models, we conducted a series of mask-filling experiments for each annotation aspect independently, using the 1000 withheld sequences from our EXP and RED datasets. We filtered the withheld sequences, retaining only those that possessed at least one annotation for the aspect under evaluation. For each sequence, we masked all annotations of the target aspect while providing the remaining annotations as context. We then assessed the models’ ability to accurately predict the masked annotations vs. ground truth with standard metrics. We evaluated four models: AT_*RED*_, AT_*EXP*_, ASM35_*RED*_, and ASM35_*EXP*_, each on their respective validation datasets. For the ASM models, we performed evaluations both with and without the corresponding protein sequences to assess the impact of sequence information on annotation prediction.

### Generation Sequence Model (GSM)

A 12 transformer block AT variant was trained on the nonredundant FINAL dataset for two epochs (AT_*FINAL*_). Then, AT_*FINAL*_ was combined with ESM2-150 in a transformer “Encoder-Decoder” scheme, where the AT last hidden state was fed to the “Decoder” through cross attention (**Figure 1D**). We modified the ESM2 model with new layer norms on the query and keys of self-attention layers to increase stability and switched the activation function to SiLU. The resulting GSM model had a protein annotation and protein sequence track, which trained both the AT and ESM2 further with MLM and cross-entropy loss, concurrently. The annotation track received annotation sequences at a set 15% masking rate. The sequence track received masked protein sequences from a noise scheduler. For the first stage of training, the masking rate was sampled from a normal distribution with a mean of 0.5, standard deviation of 0.1, and clipped at 0.15 and 1.0. The first stage received sequences with a max length of 512 for eight epochs. The annotation and sequence tracks had cross-entropy hyperparameters of 1.0 and 2.0, respectively. The first stage utilized a learning rate of 1*e*^−4^, batch size of 32, and cosine learning rate scheduler with 1000 warm-up steps. In the second stage, the mean and standard deviation were set at 0.3 and the annotation track had a mask rate of 0%. We still used a cross-entropy loss on the language modeling head output of the AT to enforce identity on the embeddings. The hyperparameters were shifted to 0.01 and 1.0 for the annotation and sequence track, respectively. During the second stage, we trained up to a max length of 2000, learning rate of 1*e*^−5^, batch size of two, gradient accumulation for an effective batch size of at least 50,000 sequence tokens, and the same learning rate scheduler and warm up. The second stage consisted of two epochs. For each epoch, the data was sampled from the Uniref50 section followed by the experimentally validated section upon completion, simulating an epoch of RED and then EXP sequentially (high and low-value tokens). The first stage used the EXP section with a maximum length of 512 and the second with the larger maximum length. For both stages, a maximum length of 256 was used for the annotation track, which encompassed all annotation examples.

### BERT-like generation

We accomplished BERT-like sequence generation by employing various popular sampling techniques, including top-*k* (considering the best *k* options per token) and nucleus sampling (thresholding and sampling options above *p*) (71). Generation from mask tokens during inference were actualized by choosing the top-*u* number of mask tokens to fill each forward pass, chosen based on the maximum logit or entropy value (before or after softmax). For instances of *k* and *p* that prevent greedy denoising, logits or entropy values were sampled from a multinomial distribution: introducing randomness into the generation process. Temperature *t <* 1.0 was used before softmax to push intra-token probabilities closer together, which heavily affects the multinomial sampling (72).

### Sequence reconstruction referencing annotations

To access the sequence reconstruction potential of dual vocabulary systems like ASM we designed a scheme to assess the effectiveness of mask filling of ASM with full annotation context versus base ESM2 models. For ease of comparison, we used *n* = 5000 and *k* = 1, which is standard greedy BERT mask filling (as *n >* sequence length). For ASM35_*EXP*_ and ASM35_*RED*_ we conducted mask filling for five percentages (5, 15, 30, 50, 70%) with five replicate experiments with different random seeds each. The same sequence sections were masked and fed to ASM or ESM2. This ensured that each model received the same sequence and mask positions, but ASM also got annotation tokens at the end. The logits were recorded, and we tracked various metrics, including F1, for tokenwise classification and cross-entropy loss.

### Sequence generation with annotation prompts

In local experiments, we observed that sequence generation with high or full masking probabilities required careful hyperparameter selection. We assessed ESM2-150 and GSM over five masking percentages (15, 30, 60, 70, 100%) over a diverse selection of *u, k* or *p*, and *t* for 100 sequences in the GSM test set.

- *u* ∈ [1, 2, 3, 10, 5000]
- *p* ∈ [0.01, 0.05, 0.10, 0.15, 0.25, 0.35, 0.45, 0.50]
- *k* ∈ [1, 2, 3, 5, 10]
- *t* ∈ [0.001, 0.01, 0.1, 0.7, 1.0, 1.5]

GSM also received the full annotations as a prompt. Of particular note, we included uncommonly low temperatures inspired by (72) in an effort to increase performance. For 100% masking, we included methionine at the start of the sequence, and tile mask tokens up until the length of the ground truth sequence.

### Multiple sequence alignment and Novel alignment score

Multiple sequence alignment was calculated with standard global alignment settings using Biopython or Biotite Python packages (73, 74). This included BLOSUM62, a gap score of −10, and gap extension of −0.5. For BLAST services, we employed SequenceServer or Blast2GO for BLASTP, for which we used the default settings and a nonredundant SwissProt reference database (54, 75, 76).

We placed multiple sequence alignment scores between zero and one by constructing an error-based metric scaled by the sequence length,

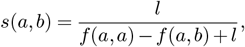

where *f* (*a, b*) is the multiple sequence alignment score using Needleman–Wunsch and BLOSUM62 between protein sequence strings *a* (ground truth) and *b* (generated sequence), and *l* is the length of string *a*. The result is an error term in the denominator that reduces the score upon poor alignment. Whereas the score scales from zero to one, we observe it is a non-linear range wherein it is increasingly difficult to get close to one. Calculated distributions of the scores can be found in **Supplemental Figure 1**.

### Low sequence identity mining

Data mining between test set and training set examples was calculated via pairwise sequence alignment and exact match accuracy for the sequence identity percentage. We filtered out low sequence identities produced by the generation repetition problem by conducting a *χ*^2^ test between the amino acid counts of a sequence vs. a reference database. The reference database reported on was the unique set of sequences from every dataset split of the GSM dataset. The null hypothesis that a sequence belonged to the reference distribution was rejected at an arbitrary *p*-value threshold 1*e*^−20^ found through manual experimentation.

## DATA AND CODE AVAILABILITY

Selected datasets, code, and model weights can be found at github.com/Gleghorn-Lab/AnnotationVocabulary.

## AUTHOR CONTRIBUTIONS

Conceptualization (LH, JPG), Annotation vocabulary (LH), Model architectures (LH), Data Curation (LH), Novel loss (LH, DH), Investigation (LH, NR, CH), Formal Analysis (LH, NR, CH, JPG), Writing – Original Draft (LH, NR, DH, JPG), Writing – Review & Editing (LH, NR, CH, DH, JPG), Supervision (JPG), Project Administration (JPG), Funding acquisition (LH, JPG).

## ACKNOWLEDGEMENTS

The authors thank Katherine M. Nelson, Ph.D., for reviewing and commenting on drafts of the manuscript. This work was partly supported by the University of Delaware Graduate College through the Unidel Distinguished Graduate Scholar Award (LH), and the National Institutes of Health through R01HL133163 (JPG) and R01HL145147 (JPG).

## Supplementary Information

## Acronyms

ASM: Annotation Sequence Model. 2, 11
AT: Annotation Transformer. 2
BERT: Bidirectional Encoder Representations from Transformers. 3
BLAST: Basic Local Alignment Search Tool. 2
BP: gene ontology Biological Process. 10
CAMP: Contrastive Annotation Modeling of Proteins. 2
CC: gene ontology Cellular Component. 10
CO: COfactor. 10
EC: Enzyme Commission number. 2
ESM: Evolutionary Scale Modeling. 2
EXP: Uniprot sequences and experimentally validated nonredundant annotations, 70,000 total. 3, 4, 10–13, 21
FINAL: Final sequence annotation dataset, built from RED and EXP principles. 10, 13
Gene3D: A database of protein domain structure annotations for protein sequences. 2
GO: Gene Ontology. 2
GSM: Generation Sequence Model. 2
LLM: Large Language Model. 2;
MCTS: Monte Carlo Tree Search. 7
MF: gene ontology Molecular Function. 10 **MLM** Masked Language Modeling. 1 **MSE** Mean Squared Error. 11
NAT: UniRef50 sequences and nonredundant natural language descriptions, 1.4 million total. 3, 4, 10
pLM: Protein Language Model. 1
PPI: Protein-Protein Interactions. 1
RED: UniRef90 sequences and nonredundant annotations, 500,000 total. 3, 4, 10–13, 21
RED_*ALL*_: UniRef90 sequences and redundant annotations, 17 million total. 10
SOTA: State-Of-The-Art. 2
TS: Thermostability. 5

---

**Table 1.**
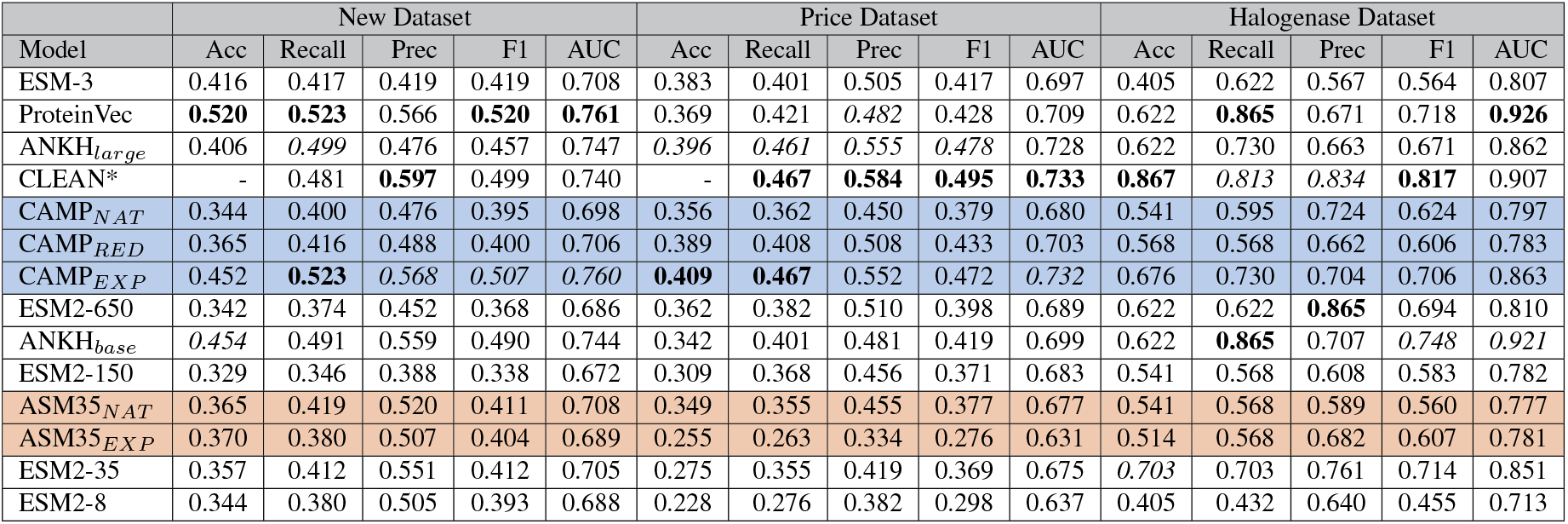
Full measured metrics for the New, Price, and Halogenase datasets via vector search. All scores except for accuracy use weighted averages, in line with the CLEAN repository. * Reported (32)

**Table 2.**
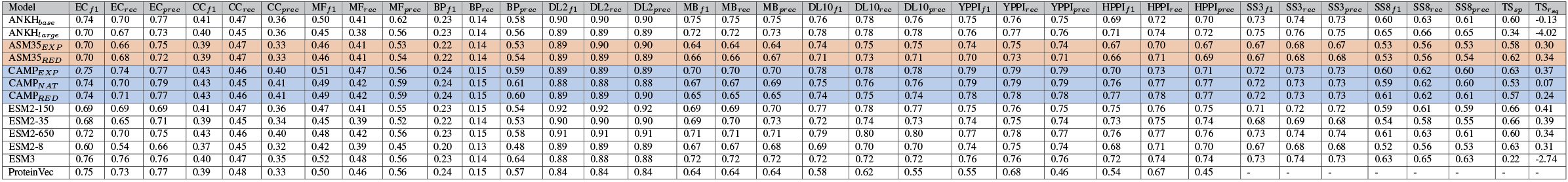
All metrics for all evaluated downstream datasets for the commonly used pLMs in the paper. Precision (prec), recall (rec), F1, accuracy (acc), Spearman *ρ* (sp), and *R*^2^ (r_*s*_*q*).

**Fig 1.**
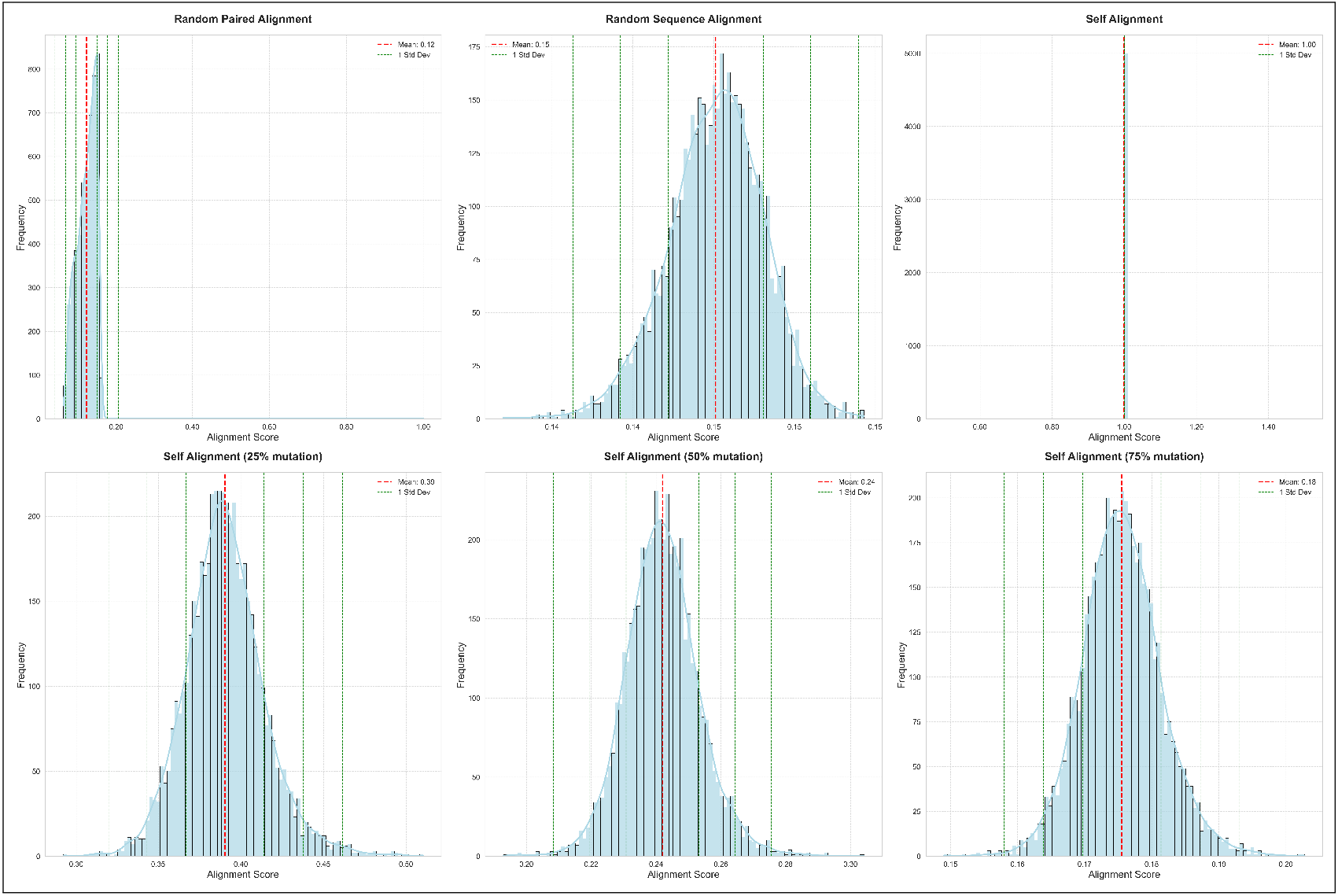
Distributions of our alignment score over various conditions, calculated over the entire GSM test set. Random paired alignment refers to random real proteins paired together. Random sequence alignment refers to a real sequence paired with a randomly generated sequence of the same length. Self alignment is alignment of a sequence with itself. 25-75% mutation refers to random mutation rates of one protein in the self alignment scheme. Overall, extremely poor sequence similarity is measured around 0.15 or lower, and good alignment is closer to one - where getting increasingly close to one requires more and more similarity.

**Fig 2.**
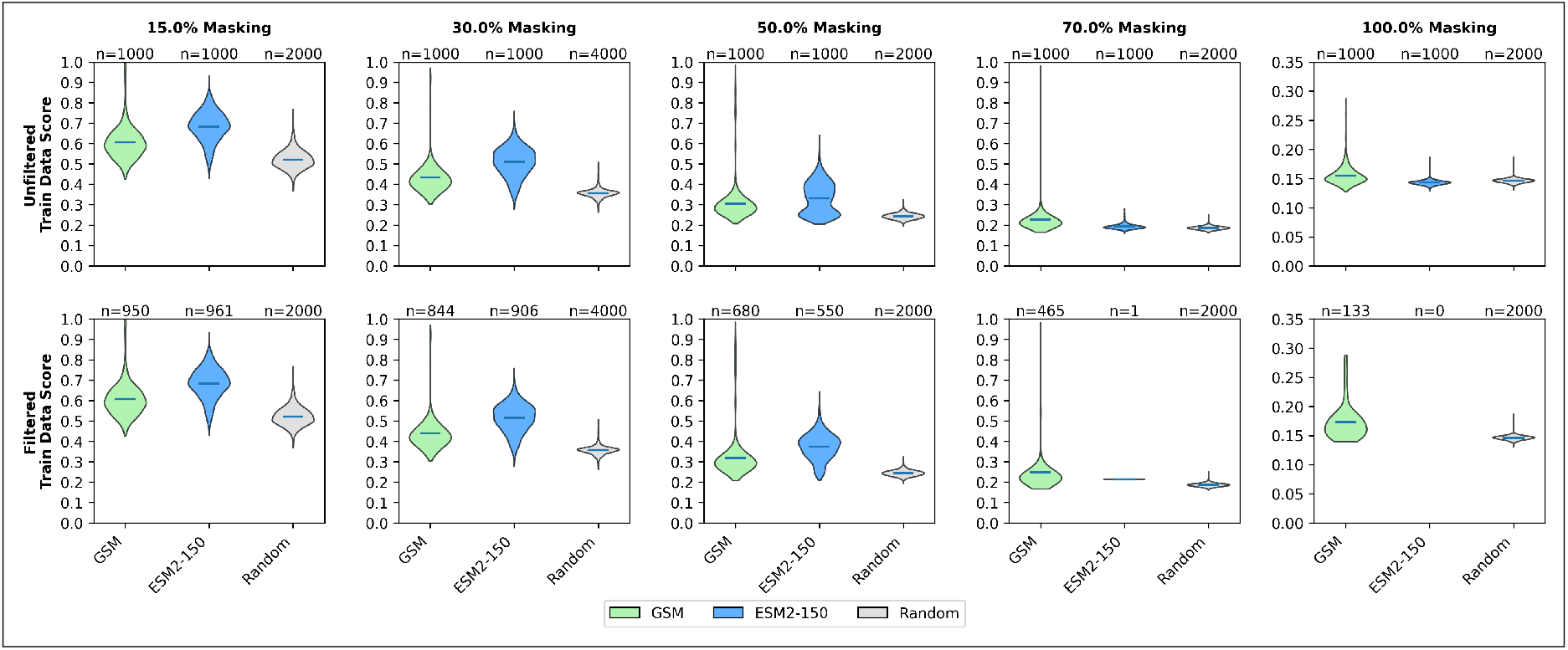
Violin plots of protein sequence generation performance over various mask parameters for 1000 random train sequences. GSM, ESM2-150, and a random mask-filling scheme receive the same masks and then fill amino acids with *u* = 10 and *k* = 1 (greedy denoising and 10 tokens at a time). However, GSM also receives an Annotation Vocabulary prompt. Low-quality results were filtered out according to amino acid distribution using *χ*2 test, labeled as “filtered.”

**Table 3.** Custom alignment score, exact match accuracy, and Levenshtein similarity for nucleus sampling *p* and temperature *t* value optimization.

|  | Parameter | Score | Accuracy | Levenshtein Similarity |
| --- | --- | --- | --- | --- |
| ESM |  |  |  |  |
| $p$ | 0.01 | 0.454 | 0.790 | 0.790 |
|  | 0.05 | 0.464 | 0.797 | 0.797 |
|  | 0.10 | 0.470 | 0.801 | 0.802 |
|  | 0.15 | 0.473 | 0.804 | 0.805 |
|  | 0.25 | 0.474 | 0.806 | 0.806 |
|  | 0.35 | **0.476** | 0.807 | 0.807 |
|  | 0.45 | 0.475 | 0.807 | 0.807 |
|  | 0.50 | 0.475 | 0.807 | 0.807 |
| $t$ | 0.001 | 0.434 | 0.609 | 0.611 |
|  | 0.01 | 0.435 | 0.608 | 0.610 |
|  | 0.1 | **0.439** | 0.607 | 0.610 |
|  | 0.7 | 0.433 | 0.604 | 0.608 |
|  | 1.0 | 0.429 | 0.601 | 0.606 |
|  | 1.5 | 0.424 | 0.599 | 0.605 |
| GSM |  |  |  |  |
| $p$ | 0.01 | 0.394 | 0.754 | 0.754 |
|  | 0.05 | 0.401 | 0.760 | 0.760 |
|  | 0.10 | 0.406 | 0.764 | 0.765 |
|  | 0.15 | 0.409 | 0.767 | 0.767 |
|  | 0.25 | **0.410** | 0.768 | 0.768 |
|  | 0.35 | **0.410** | 0.768 | 0.768 |
|  | 0.45 | **0.410** | 0.768 | 0.768 |
|  | 0.50 | **0.410** | 0.768 | 0.769 |
| $t$ | 0.001 | 0.395 | 0.601 | 0.604 |
|  | 0.01 | 0.396 | 0.604 | 0.610 |
|  | 0.1 | **0.399** | 0.611 | 0.619 |
|  | 0.7 | 0.397 | 0.606 | 0.623 |
|  | 1.0 | 0.395 | 0.604 | 0.622 |
|  | 1.5 | 0.393 | 0.601 | 0.619 |

**Table 4.** Normalized alignment scores for number of tokens per forward pass *u* and top-*k* sampling optimization. The reported score was based on alignment from the Needleman-Wunsch algorithm score (BLOSUM62) divided by the maximum sequence length: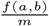, where *m* is the maximum sequence length of strings *a* and *b*.

|  | Parameter | Score (prev) |
| --- | --- | --- |
| ESM |  |  |
| $u$ | **1** | **3.488** |
|  | 2 | 3.481 |
|  | 10 | 3.463 |
|  | 5000 | 3.389 |
| $k$ | **1** | **3.484** |
|  | 2 | 3.469 |
|  | 5 | 3.447 |
|  | 10 | 3.421 |
| GSM |  |  |
| $u$ | **1** | **3.490** |
|  | 2 | 3.489 |
|  | 10 | 3.478 |
|  | 5000 | 3.328 |
| $k$ | **1** | **3.451** |
|  | 2 | 3.448 |
|  | 5 | 3.443 |
|  | 10 | 3.443 |

**Table 5.** The exact recovery accuracy of 15% random masking from the validation sets of various annotations transformers, including the AT connected to GSM with cross-attention.

|  | AT<sub>EXP</sub> | AT<sub>RED</sub> | AT<sub>FINAL</sub> | GSM |
| --- | --- | --- | --- | --- |
| Acc | 0.690 | 0.774 | 0.705 | 0.784 |

## Mixture of Experts Implementation

Mixture of Experts (MoE) extended ESM (MOESM) was our first attempt at curating multipurpose embeddings for protein annotation. We used the MoE extension method from (77) to extend pretrained ESM models with *N* identical MLP “experts” for improved performance. We made the following observations:

- Direct supervised multitask learning can lead to highly unstable loss convergence, which led to unsatisfactory results for actual annotation
- A small MOESM can use and project the embeddings of a larger pLM into functional clusters
- Token-wise MoE routing is better than sequence-wise routing, even for sequence-oriented tasks
- MOE models are highly sensitive to over-training (which has been previously observed (78)

Our best-performing MoE approach was inspired by ProteinVec, compiling similar protein pairs based on a similarity heuristic of matching EC or GO annotations, and trained with an MNR loss variant. We removed any CLEAN sequences from the dataset for downstream comparison. The model shown in **Figure 3** used frozen ANKH_*base*_ and *N* = 4 MOESM8 (from ESM8). This model also used convolutional adapters to incorporate information over all hidden states of the models, not just the last one. This model had the second (to ESM3 by 0.001 F1) best performing vector embeddings for the EC downstream task and competitive performance on CLEAN (outperforming CLEAN on the New dataset) despite its modest size of 488 million parameters (**Table 6**). We excluded from the rest of the results because of its subpar performance on average.

**Fig 3.**
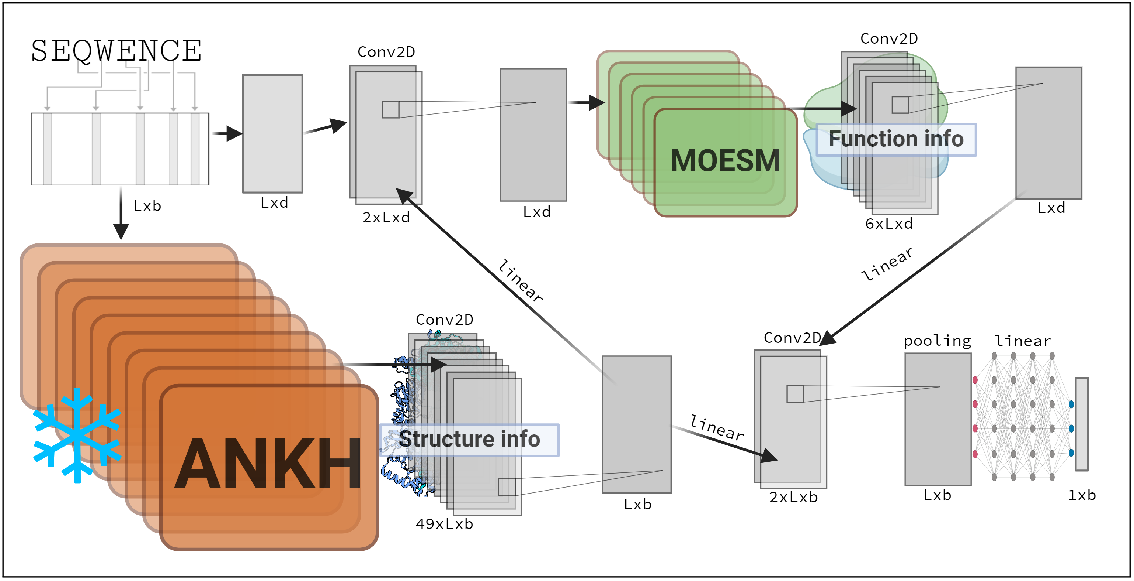
Model schematic for MOESM8. Frozen ANKH was used to build foundational numerical representations of inputs that a small MOESM added to and modified to be functionally and clustered by EC and GO. The final output was pooled and projected to a single vector for embedding large repositories. Token embedding was modified from explosion.ai. Convolution layers were made with NN-svg. Created with biorender.

**Table 6.** MOESM8 performance on downstream vector-based tasks.

| Downstream | EC | CC | MF | BP | DL2 | DL10 | Avg |
| --- | --- | --- | --- | --- | --- | --- | --- |
| F1 | 0.757 | 0.372 | 0.500 | 0.213 | 0.879 | 0.719 | 0.573 |

| CLEAN | Acc | Recall | Precision | F1 | AUC |
| --- | --- | --- | --- | --- | --- |
| New | 0.411 | 0.485 | 0.491 | 0.459 | 0.741 |
| Price | 0.369 | 0.428 | 0.477 | 0.432 | 0.712 |
| Halogenase | 0.568 | 0.757 | 0.595 | 0.634 | 0.875 |

**Fig 4.**
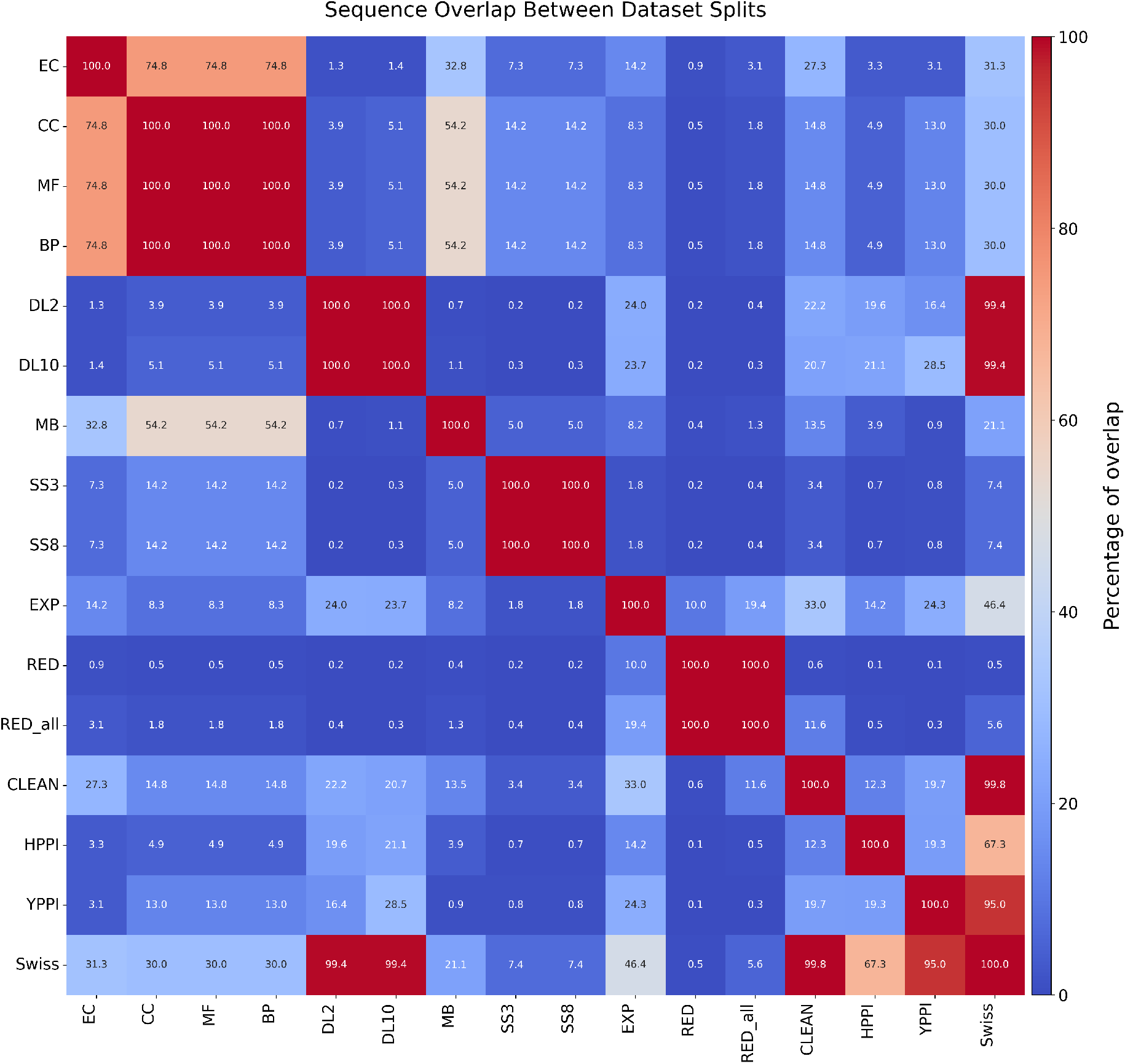
Sequence overlap by exact match between the sets of all datasets used in the paper. While our EXP dataset and RED dataset have some overlap with evaluation datasets, we are mostly interested in comparing against ProteinVec - which was trained on SwissProt (Swiss) and has much greater overlap. Additionally, because our CAMP models were trained for only one epoch, we conclude it is unlikely that this overlap biased the results.

## Full loss analysis

### Mini-batch loss

As described above, we work in each iteration with stochastic mini-batches 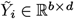 of *b* samples chosen uniformly at random (without replacement) from the full data. Namely, 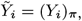_,:_ where *π* ∈ {1,…, *n*}^*b*^ denotes the *b* randomly selected sample indices (the same indices are used for all modalities). The loss evaluated on these mini-batches, i.e., 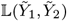, is written in code as follows:

**import** t o r c h

**import** t o r c h. nn. f u n c t i o n a l as F

**from** t o r c h m e t r i c s. f u n c t i o n a l **import** p a i r w i s e _ c o s i n e _ s i m i l a r i t y

**def** Loss (Y_1, Y_2, lambda_ 1 = 1. 0, lambda_ 2 = 0. 1): *# (b, d) (b, d)*

C_1 = p a i r w i s e _ c o s i n e _ s i m i l a r i t y (Z_1) *# (b, d)*

C_2 = p a i r w i s e _ c o s i n e _ s i m i l a r i t y (Z_2) *# (b, d)* d i f f = F. m s e _ l o s s (C_1, C_2)

a n t i _ t r i v i a l = (lambda_ 1 * C_1 + lambda_ 2 * C_2). mean ()

**return** d i f f + a n t i _ t r i v i a l

### Bias of the mini-batch loss

As described above, the mini-batch loss 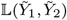 evaluated using the stochastic mini-batches 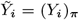 (formed from the randomly selected sample indices *π* ∈ {1,…, *n*}^*b*^) is technically biased. Specifically,

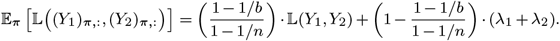

Before we derive this result, note that the bias takes the simple form of a global scaling and an added constant. The added constant has no impact on gradients, and the global scaling is straightforwardly corrected or can even be ignored by simply absorbing it into the step sizes.

We now derive this result. Noting that Θ((*Y*_i_)_*π*,:_) = (Θ(*Y*_*i*_))_*π,π*_ yields

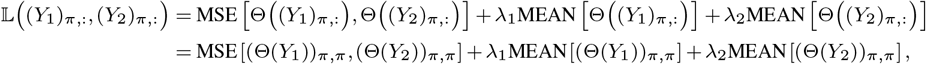

and so

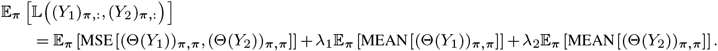

Note next that for any matrix *A* ∈ ℝ ^*n×n*^,

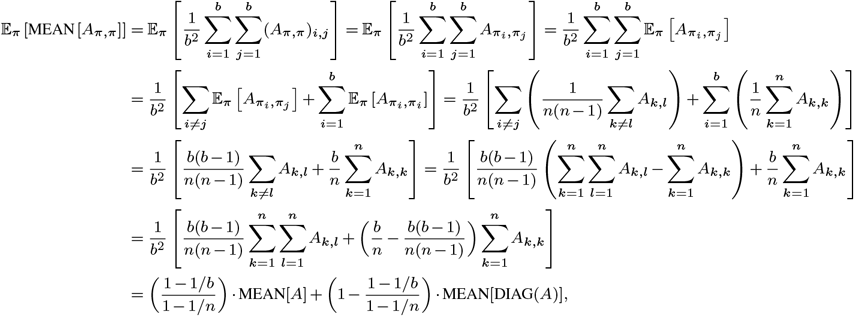

where DIAG(*A*) = [*A*_1,1_, *A*_2,2_, …, *A*_*n,n*_] ∈ ℝ ^*n*^. Thus, we have

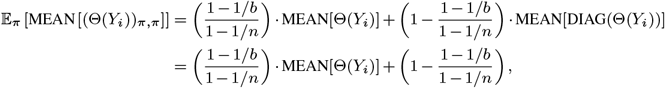

where we have used the fact that 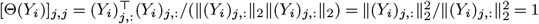. Likewise,

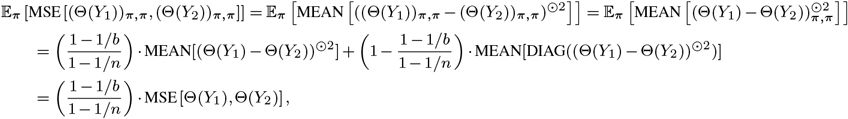

where we used the superscript ⊙2 to denote entrywise squaring. Thus, we finally have

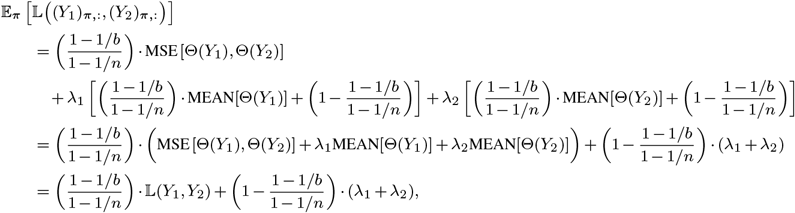

which concludes the derivation.

### Equivalent formulation of the contrastive loss

To gain further insight into the contrastive loss, this section derives an equivalent formulation. Note first that

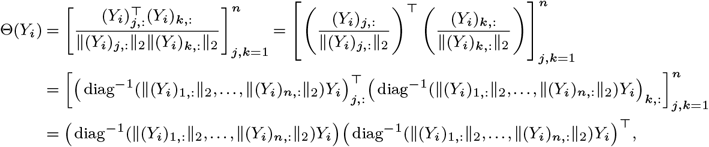

where the rows of diag^−1^(∥(*Y*_*i*_)_1,:_∥_2_, …, ∥(*Y*_*i*_)_*b*,:_∥_2_)*Y*_*i*_ are the corresponding rows of *Y*_*i*_, normalized with respect to the *𝓁*_2_ norm. As a result, the contrastive loss can also be written as

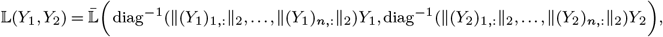

where

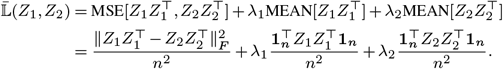

Noting that

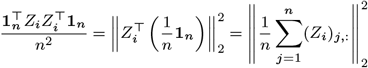

is the square norm of the average normalized row 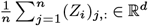 provides another interpretation of these terms. Namely, these two terms encourage the rows of *Z*_1_ and *Z*_2_ to have averages with small norms. Recall that the rows of *Z*_*i*_ all have unit norm by construction, so they are all vectors on the unit sphere in ℝ ^*d*^. Consequently, the above terms encourage these vectors to point in diverse directions (in order to induce the cancellation that would produce a small average). In other words, they encourage the rows of *Y*_*i*_ (which are vectors in ℝ ^*d*^) to point in diverse directions. However, it does not enforce that the vectors for one modality have similar magnitudes of another. The idea here is to coerce a relationship between modalities while giving models as much freedom as possible to place examples in an embedding space.

### Notes on contrastive loss performance

We assume that the regularization terms in the contrastive loss are most effective when some amount of nonredundancy is enforced upon dataset curation, so that uniform sampling produces mini-batches that do not have similar inputs *on average*. This assumption is also true for other contrastive losses such as the MNR loss, which requires minibatches to be “paired” (60). We ran small-scale experiments on scaled-down versions of CAMP using the MNR loss, the loss from cdsBERT (12), and our novel contrastive loss. MNR and our contrastive loss performed similarly on very small models, but the larger the model the more our loss showcased superior performance. However, we observe that downstream performance with our loss is *highly* sensitive to *λ*_1_ and *λ*_2_, and optimal values may be vary depending on the dataset.

